# Heterogeneous shedding and susceptibility in a *Caenorhabditis elegans* transmission model

**DOI:** 10.1101/2024.06.04.597401

**Authors:** Nic M. Vega

## Abstract

Variation in transmission plays a crucial role in shaping the dynamics of infectious diseases. Population heterogeneity is known to contribute to this variation and is often represented in epidemiological models. However, it is not always clear *a priori* what sources of variation should contribute meaningfully to a given scenario, and it can be challenging to infer distributions of underlying processes from data. In this study, we demonstrate the use of *Caenorhabditis elegans* as a tractable system in which high-quality data can be produced for experimental epidemics. We show that distributions of shedding and susceptibility in this host can be experimentally decoupled to measure heterogeneity in transmission processes. We observe and quantify super-shedding and heterogeneous susceptibility, and we show that distributions of population heterogeneity and transmission outcomes have features conserved with real-world epidemics. Our results quantify sources of heterogeneity in bacterial transmission in this small model organism and establish *C. elegans* as a promising quantitative model for experimental epidemics.

## Introduction

Variation in transmission is well known to contribute to the population-level behavior of infectious diseases. The number of secondary infections caused by an individual case can vary dramatically, with some individuals and/or events causing large numbers of secondary infections and others none. When population size and incidence of infection are sufficiently high, the erratic contributions of individuals will be somewhat smoothed by large number effects (Grassly and Fraser 2008; Hébert-Dufresne et al. 2020). However, stochasticity and heterogeneity strongly affect transmission, and the resulting variation in transmission shapes the range of possible outcomes for an outbreak (Rose et al. 2021).

In this work, we define these terms according to common practice in theoretical ecology: *heterogeneity* refers to differences in demographic traits between individuals, and/or in the distributions of these traits across populations, and *stochasticity* refers to internal (demographic) or external (environmental) noise (Kendall and Fox 2003). We use *variation* as an agnostic term to describe distributions of outcomes as observed from data. In distributions of observed events, we can describe variation by calculating *sample variance;* if there are named distributions fitted to these observations, we can estimate *variance* as a parameter. In this context, outbreaks are inherently stochastic – early transmission involves small numbers of events, and process-based or demographic stochasticity can create very different trajectories for epidemics with identical parameters (Grassly and Fraser 2008). Further, as described above, transmission is highly variable; both heterogeneity and stochasticity can contribute to this variation (Galvani and May 2005; Woolhouse et al. 1997).

In terms of model structure, heterogeneity is generally considered as a major source of variation. The processes that contribute to transmission heterogeneity differ across outbreaks, with consequences for epidemic outcomes (Rose et al. 2021; Gomes et al. 2022; Bansal, Grenfell, and Meyers 2007; Britton, Ball, and Trapman 2020; Hébert-Dufresne et al. 2020). Some of the relevant factors are biological (e.g. host genetics and life history; lineage of the transmitted agent). Others depend on behavior and the physical and social environment – for example, the number and/or intensity of connections among individuals. Sources of heterogeneity can differ across outbreaks, with consequences for R0 (Woolhouse et al. 1997) and efficacy of control measures (Lloyd-Smith et al. 2005; Sneppen et al. 2021).

Heterogeneity in epidemic models is inferred by fitting models to data. However, data from real-world epidemics are frequently complicated in ways that can make models difficult to specify and biases in inference difficult to assess (Gostic et al. 2020). Real-world data have common flaws (sparse/under-sampled, with uneven effort, in populations where most heterogeneity is hidden) and simulated data are limited by our understanding of real data. It is therefore difficult to determine the form of variation in shedding and/or susceptibility within and across populations, or to understand how different sources of variation affect distributions of transmission.

Small host models provide opportunities to bridge the gap by generating high-quality data in controlled populations. Experimental work in insects and other small animal hosts has demonstrated the potential of experiments for advancing epidemic modeling (Knell, Begon, and Thompson 1998; D’Amico et al. 1996; Dwyer and Elkinton 1993; Bouma, de Jong, and Kimman 1995; Reeson et al. 2000; Barlow 2000; Dobson and Meagher 1996; Ben-Ami, Regoes, and Ebert 2008). However, extant small host systems mostly represent scenarios of specific interest, where a disease of economic or environmental concern exists in a host where experiments happen to be feasible. These non-model hosts are in general slow to grow, limited in numbers, and difficult to monitor with high resolution. For this reason, experimental studies of transmission in animal models are few and typically contain data from small numbers of infected individuals and/or transmission events, for example (Knell, Begon, and Thompson 1998; Greer, Briggs, and Collins 2008; Sadd and Barribeau 2013; Neri et al. 2011; Jankowski et al. 2013; Tadiri, Fussmann, and Scott 2021). A tractable, quantitative, high-throughput small animal model with the biological capacity to simulate realistic epidemics would allow direct tests of theory and the possibility of learning new rules of transmission.

Here we use a tractable model organism, the nematode *Caenorhabditis elegans*, as a platform for experimental epidemics. *C. elegans* provides large, well-characterized, repeatable populations wherein excellent control of colonizer, host, and environment can be achieved. Illustrating its biological relevance, the worm is a well-established system for studying host-microbe interactions (Berg et al. 2019; Rafaluk-Mohr et al. 2018; Dirksen et al. 2016; Shapira 2017) in large populations where variation can be measured accurately (Taylor, Spandana Boddu, and Vega 2022). The ability to generate large, repeatable populations of hosts is critical for observing rare events and describing distributions of outcomes.

We demonstrate quantification of transmission heterogeneity from experimental data in the *C. elegans* model. We show how shedding and secondary infections can easily be decoupled in this system, allowing these distributions to be described and manipulated separately. We replicate transmission events across different susceptible populations to describe distributions of secondary infections as coupled functions of shedding and susceptibility. We show that, as in real-world epidemics, most infected individuals produce no secondary infections and a small fraction of hosts are responsible for the majority of transmission. Further, we find that variation in shedding and susceptibility is not well described by simple models, with consequences for predicted distributions of secondary infections. Our results quantify sources of heterogeneity in bacterial transmission by *C. elegans* and demonstrate the potential utility of this host as a novel high-throughput system for experimental epidemics.

## Results

### (Super)-shedding in C. elegans

Super-spreading is common in transmitted diseases, with a commonly referenced “80/20 rule”, wherein roughly 80% of transmission is attributable to 20% of cases. The observed ratio can and does vary, but the general pattern emerges empirically from data across a wide range of pathogens (Galvani and May 2005; Woolhouse et al. 1997; Lloyd-Smith et al. 2005). Variation in transmission can be attributed to biological super-shedding, differences in contacts or behavior, or combinations of factors (May and Anderson 1987; Endo et al. 2020; Susswein and Bansal 2020; Galvani and May 2005; Chowell et al. 2016). The distinction is not trivial - different forms of super-spreading lead to qualitatively different distributions for transmission (Kuylen et al. 2022).

We therefore first sought to describe shedding in *C. elegans*, including prevalence of super-shedding, and to understand the factors affecting shedding rates. In our experiments, live fluorescently labeled bacteria are shed from the intestines of pre-colonized (index case) worms into the surrounding liquid medium, where growth of bacteria can be measured. This allowed us to observe shedding by individual worms without additional variation due to transmission into new hosts.

For these and later experiments, the primary infectious agent used was the Gram-negative pathogen *Salmonella enterica* LT2 bearing a chromosomally integrated GFP marker. *S. enterica* is a slow-killing intestinal pathogen of *C. elegans* (Aballay et al. 2003; Aballay, Yorgey, and Ausubel 2000). There is a delay of several days post-infection before *S. enterica*-induced morbidity/mortality is observed (Biancalani and Gore 2019); on the time scale of our experiments, host death is essentially zero and can be neglected. Additionally, to demonstrate how shedding can differ across infectious agents, we measured shedding of the Gram-positive pathogen *Staphylococcus aureus* Newman (Begun et al. 2005; Sifri et al. 2003), and the native worm microbiome commensal *Ochrobactrum* MYb14 (Dirksen et al. 2016; Zimmermann et al. 2020). All bacterial strains used are strong colonizers, reaching saturation in the gut (peak 10^5^-10^6^ bacteria/worm) within 24 to 48 hours (**Figure S1**). For the pathogens *S. enterica* and *S. aureus*, immunity and pathogenesis in *C. elegans* are described (Sifri, Begun, and Ausubel 2005; Garsin et al. 2003). The commensal MYb14 colonizes the gut to comparable densities without increasing mortality and may benefit the host (Zimmermann et al. 2020). When worms are colonized with labeled bacteria, fluorescence of individual worms increases monotonically with live bacterial load and is a useful proxy for colonization (Murfin, Chaston, and Goodrich-Blair 2012; Twumasi-Boateng, Berg, and Shapira 2014), with the caveats that (1) thresholds of detection exist and (2) fluorescence can saturate for highly colonized worms, depending on the intensity of the reporter (**Figure S1**).

Briefly, in these experiments, N2 adult worms pre-colonized with each bacterial agent (henceforth “index cases”) were sorted into bins based on GFP fluorescence in individual worms (**Figure S1**). Worms were then chilled to halt feeding and excretion, lightly bleached to remove external bacteria, then transferred individually into wells of a 96-well plate containing S medium + heat-killed OP50 as an inert food source to allow shedding. Adult worms recover rapidly from short-term cold paralysis, allowing normal activity – including excretion cycles and shedding – to resume within 5-15 minutes after returning to normal temperature (Robinson and Powell 2016; Manjarrez and Mailler 2020). For simplicity, the time at which index case worms were transferred to individual wells was used as time 0 of these experiments.

These index cases shed live bacteria from the intestine into the liquid supernatant, where the bacteria proliferate, and shedding was measured by plating small samples of the supernatant over time (**Figure 1A**). The waiting time before a countable density is achieved depends on both the waiting time for a shedding event and the rate of bacterial growth in the supernatant. Samples of supernatant were therefore plated at specified intervals to monitor for appearance of live bacteria, and bacteria were considered “countable” at a minimum of 4 colonies/spot (**Methods**). Using the first data point from each well at which a countable number of colonies were observed, the time at which the first shedding event occurred in that well (henceforth *WaitTime*) was calculated by assuming that growth at these very low densities was exponential (*N*(*t*) = *N*_0_*e*^*rt*^), using values for bacterial growth rate *r* calculated within each experimental run. As shedding is commonly modeled as a Poisson process with rate λ, for which the waiting times are exponential with mean 1/ λ, the average of *WaitTime* across worms within a data set therefore provided an estimator for the rate parameter. In some wells, the number of bacteria at the first countable time point was higher than could be accounted for with these assumptions, resulting in a negative (invalid) estimate for *WaitTime*; these worms were labeled as “super-shedders” and were not included in the exponential rate calculation. Note that “super-shedders” were labeled based on magnitude of the *first* successful shedding event, as the contributions of subsequent shedding events will be swamped by exponential growth of bacteria from the first event when shedding is rare. Wells not confirmed to contain a live, intact worm at the experiment’s end were removed from analysis.

**Figure 1.**
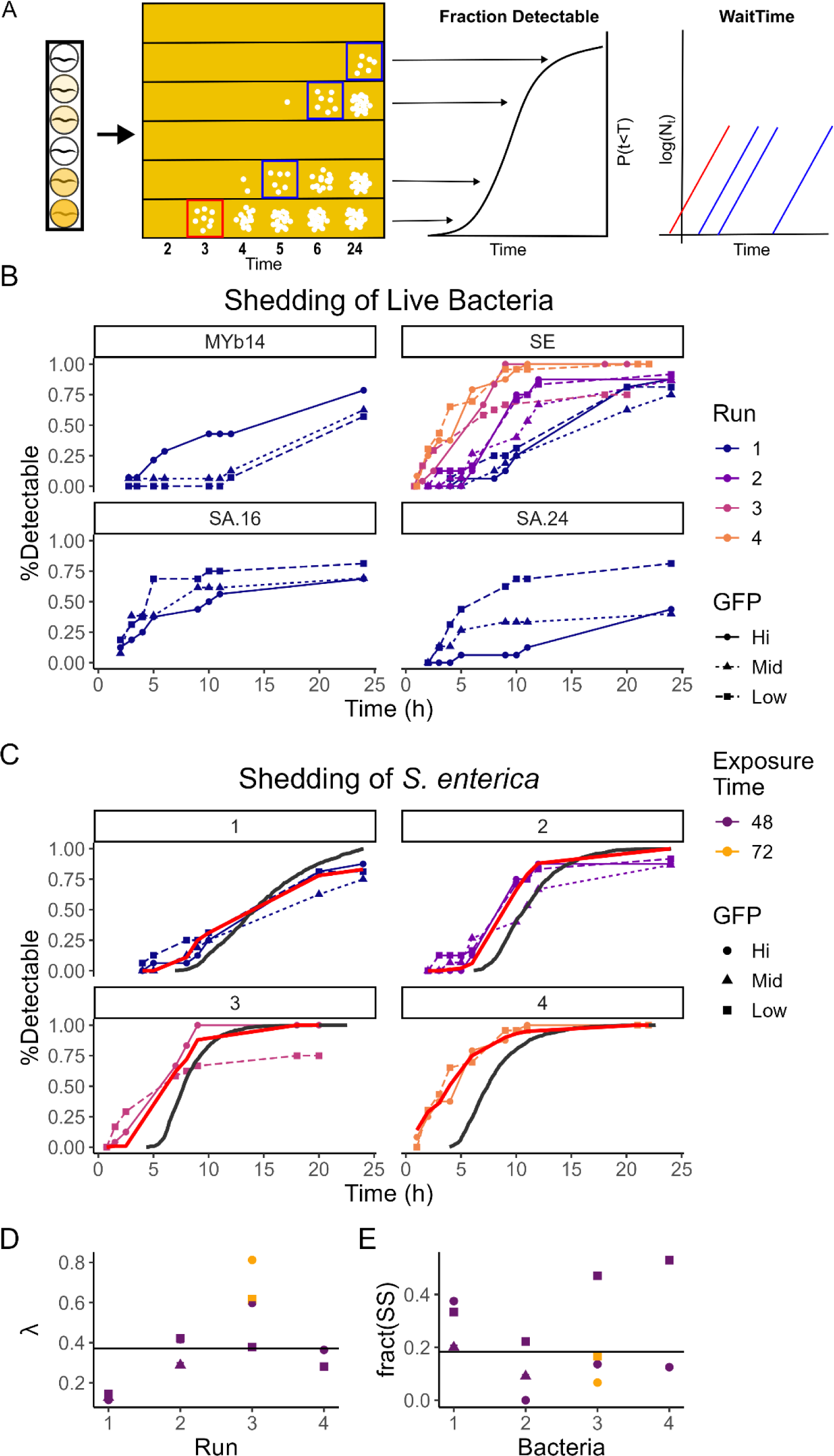
Shedding of live bacteria from the intestine of *C. elegans.* (A) Experimental schematic for shedding experiments. Index case worms were separated into individual wells of a 96-well plate, and bacterial shedding was monitored by plating aliquots of supernatant from individual wells to determine CFU/well and cumulative fraction of wells with detectable shedding (Fraction Detectable) over time. Time points where a countable number of colonies were detected (blue boxes) were used to back-calculate the time of the first shedding event in each well, *WaitTime*, assuming exponential growth from an initial shedding event. Wells where this calculation generated a negative value for the time of the first shedding event (red) were considered “super-shedders” and were not included in distributions of *WaitTime*. (B-C) Fraction of wells with detectable bacteria over time. Independent experimental runs are shown with different colors; GFP bin to which index cases belonged is indicated by point shape and line type. (B) Shedding varies across bacteria, with different dependences on bacterial load and time since exposure. Data are fraction of wells (8-24 wells per condition) above the detection threshold (1 colony/µL undiluted supernatant), measured over time during shedding by individual N2 worms into liquid media in a 96-well plate. (C) Shedding of *S. enterica* is consistently under-estimated by the waiting-time distribution for individual worms. Dark grey solid lines represent Gillespie simulations of cumulative fraction of wells with detectable shedding, using exponential waiting-time rates estimated from non-super-shedder worms. Red lines represent GSSA simulations with the same waiting-time distributions and Pareto shedding. (C-D) Parameter estimates for shedding of *S. enterica*. Estimated values for (C) λ (rate constant) and (D) fraction super-shedders (frac(SS)) across GFP bins and independent runs of the experiment. Black lines represent median values.

Overall, shedding of live bacteria occurred at low rates (**Figure 1B-D**), with estimated Poisson rate parameters ranging from ∼0.07-0.7 bacteria*h^-1^ (**Table 1**). The per-worm bacterial load and the time since initial exposure sometimes affected shedding, but the impact of each factor varied across infecting bacteria. The naive assumption that higher bacterial load leads to higher shedding held true only for the commensal MYb14; this reflected a change in the fraction of super-shedders rather than in the inferred shedding rate (**Table 1**). For the two pathogens, we observed either no relationship (*S. enterica; S. aureus* 16 hours post-exposure) or a negative relationship (*S. aureus* 24 hours post-exposure) between bacterial load and shedding. Further, for *S. aureus* but not *S. enterica*, increasing time since initial exposure appeared to decrease shedding (**Figure 1B**), possibly due to the more rapid onset of morbidity/mortality after *S. aureus* infection.

**Table 1.** Summary of shedding by *C. elegans*. Summary statistics are shown for index cases colonized with each bacteria (MYb14*, S. enterica, S. aureus*) by run and GFP bin. *Worm Age* is the age in day of adulthood of worms at the start of the experiment. *InfecTime* is the duration of exposure to GFP-labeled bacteria used to create index cases. *n* is the total number of index case worms in each data set. *nDetect* and *fDetect* are the number and fraction of individual worms with detectable shedding at 24 hours; *nCount* is the number of worms with a countable number of colonies at any time point. *λ* is the estimated shedding rate parameter for each experiment; *SE λ* is its jackknife standard error. *nSS* and *fSS* are the number and fraction of super-shedders in each data set; *SE_fSS* is the jackknife standard error for fraction super-shedders. Note that the extended lag time of *S. aureus* prevented inference of shedding rates and super-shedder frequency.

| Bacteria | Run | Worm Age | Infect Time | Bin | n | nDetect | fDetect | nCount | $\lambda$ | $SE_{\lambda}$ | nSS | fSS | $SE_{fSS}$ |
| --- | --- | --- | --- | --- | --- | --- | --- | --- | --- | --- | --- | --- | --- |
| MYb14 | 1 | 1 | 48 | Hi | 14 | 11 | 0.79 | 10 | 0.120 | 2.30E-03 | 5 | 0.50 | 1.22E-02 |
| MYb14 | 1 | 1 | 48 | Low | 14 | 8 | 0.57 | 7 | 0.118 | 1.39E-03 | 0 | 0.00 | 0.00E+00 |
| MYb14 | 1 | 1 | 48 | Mid | 16 | 10 | 0.63 | 10 | 0.097 | 4.58E-04 | 2 | 0.20 | 9.07E-03 |
| *S. enterica* | 1 | 2 | 48 | Hi | 16 | 14 | 0.88 | 8 | 0.115 | 1.50E-03 | 3 | 0.38 | 7.62E-03 |
| *S. enterica* | 1 | 2 | 48 | Low | 16 | 13 | 0.81 | 9 | 0.145 | 2.87E-03 | 3 | 0.33 | 8.17E-03 |
| *S. enterica* | 1 | 2 | 48 | Mid | 16 | 12 | 0.75 | 10 | 0.126 | 1.89E-03 | 2 | 0.20 | 7.58E-03 |
| *S. enterica* | 2 | 2 | 48 | Hi | 8 | 7 | 0.88 | 7 | 0.418 | 1.62E-02 | 0 | 0.00 | 0.00E+00 |
| *S. enterica* | 2 | 2 | 48 | Low | 24 | 22 | 0.92 | 18 | 0.422 | 3.24E-03 | 4 | 0.22 | 3.67E-03 |
| *S. enterica* | 2 | 2 | 48 | Mid | 15 | 13 | 0.87 | 11 | 0.288 | 1.11E-02 | 1 | 0.09 | 5.52E-03 |
| *S. enterica* | 3 | 2 | 48 | Hi | 24 | 24 | 1.00 | 22 | 0.598 | 4.71E-03 | 3 | 0.14 | 3.00E-03 |
| *S. enterica* | 3 | 2 | 48 | Low | 24 | 18 | 0.75 | 17 | 0.378 | 9.02E-03 | 8 | 0.47 | 5.28E-03 |
| *S. enterica* | 3 | 2 | 72 | Hi | 24 | 24 | 1.00 | 15 | 0.812 | 4.96E-03 | 1 | 0.07 | 1.81E-03 |
| *S. enterica* | 3 | 2 | 72 | Low | 24 | 23 | 0.96 | 18 | 0.616 | 4.70E-03 | 3 | 0.17 | 3.12E-03 |
| *S. enterica* | 4 | 1 | 48 | Hi | 24 | 24 | 1.00 | 16 | 0.364 | 3.62E-03 | 2 | 0.13 | 2.51E-03 |
| *S. enterica* | 4 | 1 | 48 | Low | 23 | 23 | 1.00 | 17 | 0.281 | 2.06E-03 | 9 | 0.53 | 4.73E-03 |
| *S. aureus* | 1 | 1 | 16 | Hi | 16 | 11 | 0.69 | 11 |  |  |  |  |  |
| *S. aureus* | 1 | 1 | 16 | Low | 16 | 13 | 0.81 | 12 |  |  |  |  |  |
| *S. aureus* | 1 | 1 | 16 | Mid | 13 | 9 | 0.69 | 6 |  |  |  |  |  |
| *S. aureus* | 1 | 1 | 24 | Hi | 16 | 7 | 0.44 | 6 |  |  |  |  |  |
| *S. aureus* | 1 | 1 | 24 | Low | 16 | 13 | 0.81 | 11 |  |  |  |  |  |
| *S. aureus* | 1 | 1 | 24 | Mid | 15 | 6 | 0.40 | 2 |  |  |  |  |  |

The focus of these experiments was on shedding of *Salmonella enterica* (**Figure 1C-E**) due to this agent’s long history as a pathogen of *C. elegans.* Duration of exposure to *S. enterica* (72 vs 48 hours, run 3) did not affect bacterial load (**Figure S1E**) but slightly increased point estimates of shedding rates (**Figure 1C**). Inferred shedding rate varied between runs of the experiment, but most estimates were ∼0.4 bacteria/h (0.3-0.6) (**Figure 1D**), and bacterial load in the intestine had no significant effect on shedding (**Figure S2A**). The observed fraction of super-shedders likewise varied across runs but was generally 10-30%, consistent with the expected “80/20” rule. Low-GFP individuals colonized with *S. enterica* were more likely to super-shed overall (binomial GLM formula *isSuperShedder ∼ Bin*, BinLow = +1.1, p = 6.2e^-3^, df = 235)(**Figure 1E**, **Table 1**), consistent with prior work in a fish-parasite system (Stephenson et al. 2017), but final bacterial load in individual worms was not significantly associated with super-shedder status (binomial GLM formula *isSuperShedder ∼ logCFU +Bin*, logCFU = -+0.25, p = 0.51, df = 25) (**Figure S2B-C**).

The shedding rates derived from waiting-time estimates consistently under-estimated total shedding (**Figure 1C**), confirming a straightforward expectation - the population-based average shedding rate will be biased upwards due to super-shedders. We therefore modeled super-shedding using an approach where the number of events in a period remains Poisson but the size of the events follows a Pareto distribution (Pickands III 1975; Balkema and Haan 1974), as has been done recently to describe SARS-CoV-2 super-spreading (Wong and Collins 2020). In this approach, magnitude of shedding was described as Pareto, with the number of live bacteria shed in a single event distributed as 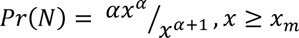, where *x_m_*is the location and α is the scale of the Pareto process. The waiting time for shedding events was assumed to be exponentially distributed as before, using rate parameters estimated from non-super-shedding worms. We obtained estimates for α by fitting GSSA predictions to data for fraction detectable wells over time, with all other parameters held constant within a run. Estimates for α were very consistent over the first three runs (α=1.3-1.7) but run 4 gave a different estimate (α=0.23) due to early detection of bacteria in some wells of this experiment. The addition of Pareto super-shedding was sufficient to account for observed distributions of bacterial shedding (**Figure 1C**). As a check, we also inferred the number of bacteria required to explain colony counts assuming that the first shedding event occurred at time 0; these inferred shedding event distributions were likewise consistent with a Pareto distribution (**Figure S3**), with shape estimates in [0.7, 1.3] and slightly higher α estimates in [2.2, 2.8] except for run 1 with α = 4.8 ± 2.4.

### Heterogeneous susceptibility

During infection, initially susceptible hosts transition to the infected state following exposure to the agent. A classical SI model with homogeneous susceptibility and constant host population size gives a Poisson distribution of secondary infections and a linear relationship between log(fraction infected) and time, −*log*(*S*(*t*)⁄*S*(0)) = β*Pt*, where S(0) and S(t) are the size of the uninfected (susceptible) population at time 0 and at time *t*, *β* is a rate constant for infection, and *P* represents intensity of exposure to the infectious agent (e.g. encounters with infected individuals or agent density in the environment). In a heterogeneous population, where susceptibility has a distribution across the population rather than a constant value, the most susceptible individuals will be infected early and/or at low exposures (Rose et al. 2021; Dwyer, Elkinton, and Buonaccorsi 1997), decreasing average susceptibility in the remaining un-infected population. For example, when susceptibility is a Gamma r.v. with dispersion *k* and average susceptibility 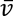, the distribution of secondary infections becomes negative binomial, and accumulation of new infections over time is sub-linear, 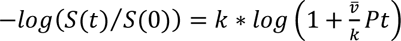 (Dwyer, Elkinton, and Buonaccorsi 1997). (While the Gamma distribution provides this specific relationship, sub-linearity holds if susceptibility is heterogeneous regardless of the distribution of β.) Parameters can be estimated directly by fitting models to dose-infectivity data, where initially susceptible populations are exposed to a range of infectious titers and/or durations (Ben-Ami, Regoes, and Ebert 2008; Dwyer, Elkinton, and Buonaccorsi 1997; Orlofske et al. 2018; Rachowicz and Briggs 2007; Barlow 2000; D’Amico et al. 1996; Dwyer and Elkinton 1993; Ben-Ami, Ebert, and Regoes 2010). Susceptible populations can differ in susceptibility due to differences in genetics, life history, etc, resulting in different parameterizations for these models (White, Forester, and Craft 2018; Galvani and May 2005).

Across simple models with and without heterogeneity, there is a common implicit assumption that accumulation of new infections is a function of exposure over time, expressed here via a cumulative term of the form *Pt* (exposure dose or intensity * time). In a stochastic model, this is interpretable as a statement about probability of infection per encounter. This is conceptually useful, but the interaction between exposure intensity and time need not be as simple as implied by the cumulative term. For example, infectious agents in general have minimal infectious doses below which infection will not occur; when this is relevant, a continuous model will produce more infections than reality.

To determine whether susceptibility in *C. elegans* is heterogeneous, and to clarify the contributions of infectious dose vs exposure duration on infection rates, we exposed initially germ-free populations of adult worms (1-6 replicate wells per experiment, 30-100 worms/well) to fixed concentrations of fluorescently labeled bacteria (10^7^-10^10^ CFU/mL). After an initial exposure of pre-determined duration (2-24 hours), the infectious agent was removed, and worms were moved to inert food + antibiotic to allow out-growth of any initial infections while preventing re-inoculations. Successful infections were identified based on fluorescence after 24-48 hours of out-growth, with a threshold for infection set at the 95^th^ percentile of fluorescence in un-infected controls (no exposure to fluorescent bacteria). Two host genotypes (N2 wild type and *daf-16* “immune-compromised”) were used to demonstrate how host genotype can affect distributions of susceptibility. The pathogen *S. aureus*-GFP was not used in these experiments to avoid the need to consider death of infected individuals, and because GFP threshold of detection for this strain was high (**Figure S2)**.

Accumulation of new infections differed between host lineages and infectious agents (**Figure 2, Table 2**). For the commensal MYb14, N2 was generally slightly more susceptible than *daf-16* (slightly higher 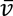). By contrast, for the pathogen *S. enterica*, N2 had lower susceptibility than *daf-16*. For both agents, *daf-16* was less heterogeneous in susceptibility than N2 (higher *k*, Gamma model). Incubation time on the inert food source was not positively correlated with either infection prevalence or intensity, indicating a lack of re-inoculation as expected (**Figure S4**). However, we observed a negative correlation between infection prevalence and incubation time for *S. enterica* (Spearman correlations: N2, rho=-0.59 p=1.3e-12, and *daf-16*, rho=-0.63 p=2.27e-7; MYb14, all Spearman rho p>0.05), suggesting some recovery for infections by *S. enterica* but not the commensal MYb14.

**Figure 2.**
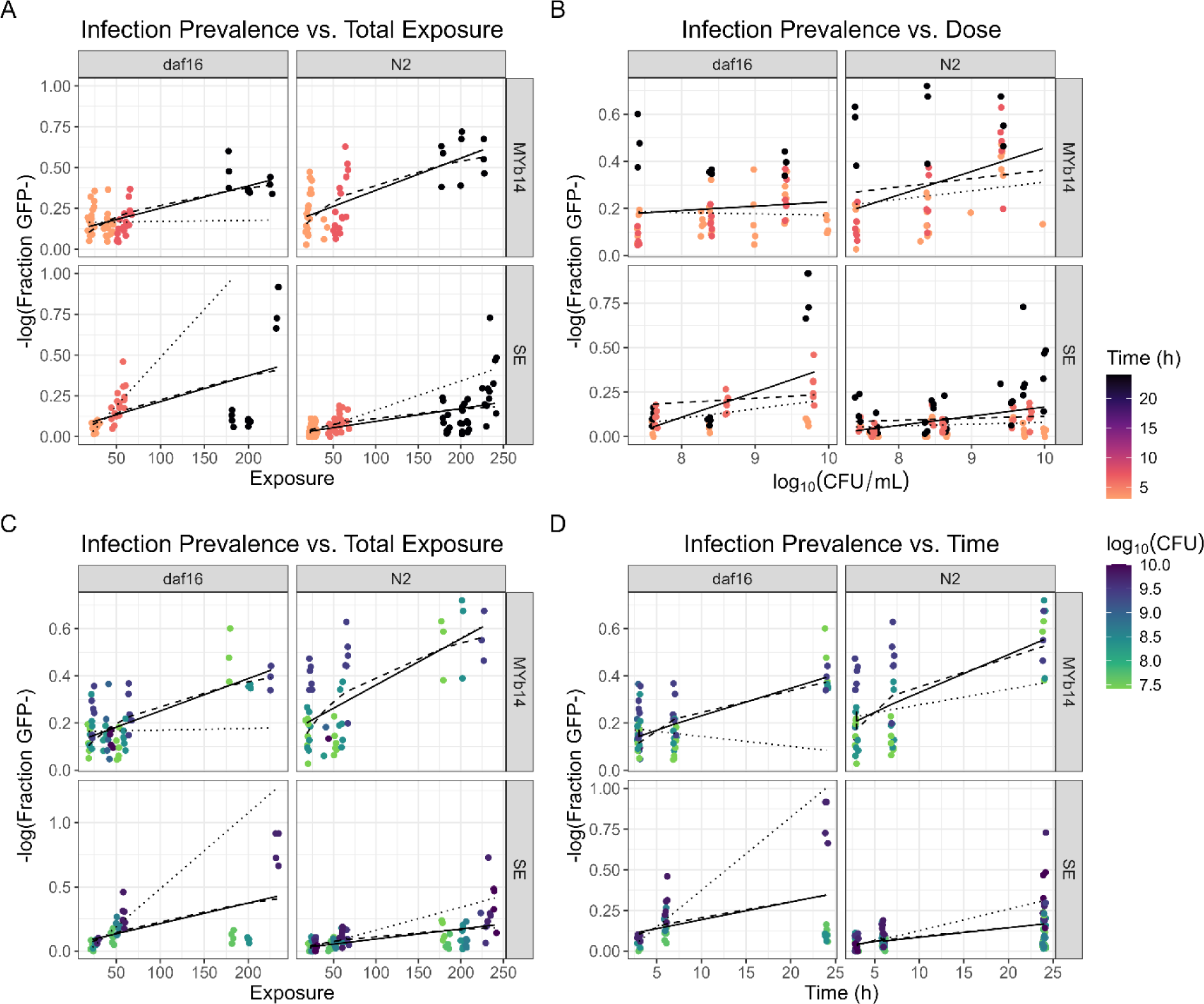
Susceptibility of N2 and *daf-16* hosts in liquid media. “Time” is the duration of exposure to the agent in hours. “Exposure” is the product of log10(bacterial density in media)*exposure time. Data are fraction of individual hosts per well that are above threshold of detection for GFP fluorescence, defined as 95th percentile total GFP/worm in un-colonized control worms (1-12 replicate wells per experiment, 50-80 worms/well). Color indicates (A-B) duration of exposure (hours) or (C-D) bacterial concentration in media (CFU/mL). Note that (A,C) are the same plot with data colored by either (A) exposure time or (C) dose. After exposure, worms were moved to inert food (heat-killed OP50) + chloramphenicol to allow growth of intestinal bacteria while preventing new infections. Lines show fits of the linear (solid black line) and Gamma (dashed black line) susceptibility models to each full data set, as well as fits of the linear model to low-exposure data (dotted line; Exposure<100, time<=7, log10CFU<9).

**Table 2.** Regression parameters for fits of homogeneous (SI) and heterogeneous (Gamma) susceptibility models. In all cases, the dependent variable was *-ln(S(t)/S0)*; the independent variable is shown for each set of parameter estimates. Linear models were fit using *lm()* and nonlinear models using *nls()* in R, with defaults unless otherwise stated. For *S. enterica*, fits were weighted using number of worms per sample; weighting had minimal effect on MYb14 fits, and so unweighted fits are shown. Values in red indicate data sets where *nls()* failed to converge; estimates are from *optimize()* with initial values (0.2, 0.02), lower bounds (0.01, 0.001), and upper bounds (10, 1). Significance codes: ***, p<0.001; **, p<0.01; *, p<0.05;., p<0.1.

| Cumulative exposure (log(CFU)*time) |  |  |  |  |  |  |  |  |
| --- | --- | --- | --- | --- | --- | --- | --- | --- |
| Host | Bacteria | Gamma susceptibility |  |  | Homogeneous susceptibility |  |  | df |
|  |  | k | vbar | RSE | m | b | RSE |  |
| N2 | MYb14 | 0.22*** | 0.01*** | 0.15 | 2e-4*** | 0.17*** | 0.15 | 46 |
| *daf-16* | MYb14 | 0.16*** | 7.9e-3*** | 0.10 | 1.3e-3*** | 0.11*** | 0.09 | 58 |
| N2 | *S. enterica* | 0.21 | 1.4e-3** | 0.82 | 7.6e-4*** | 2.3e-2 | 0.81 | 94 |
| *daf-16* | *S. enterica* | 0.25 | 3.7e-3 . | 1.44 | 1.5e-3*** | 5.3e-2 | 1.42 | 40 |

| Exposure time only |  |  |  |  |  |  |  |  |
| --- | --- | --- | --- | --- | --- | --- | --- | --- |
| Host | Bacteria | Gamma susceptibility |  |  | Homogeneous susceptibility |  |  | df |
|  |  | k | vbar | RSE | m | b | RSE |  |
| N2 | MYb14 | 0.195** | 0.11*** | 0.16 | 0.17*** | 0.016*** | 0.15 | 46 |
| *daf-16* | MYb14 | 0.15*** | 0.07*** | 0.10 | 0.11*** | 0.012*** | 0.09 | 58 |
| N2 | *S. enterica* | 0.1* | 0.02* | 0.85 | 5.9e-3 | 0.032 | 0.85 | 94 |
| *daf-16* | *S. enterica* | 0.14 | 0.05 | 1.50 | 9.8e-3** | 0.082 . | 1.52 | 40 |

| Bacterial density only |  |  |  |  |  |  |  |  |
| --- | --- | --- | --- | --- | --- | --- | --- | --- |
| Host | Bacteria | Gamma susceptibility |  |  | Homogeneous susceptibility |  |  | df |
|  |  | k | vbar | RSE | m | b | RSE |  |
| N2 | MYb14 | 10 | 3.7e-2 |  | 0.1** | -0.54* | 0.18 | 46 |
| *daf-16* | MYb14 | 0.37 | 3.0e-2 | 0.12 | 4.4e-2 | 1.8e-2 | 0.12 | 58 |
| N2 | *S. enterica* | 10 | 1.2e-2 |  | 3.9e-2** | -0.23* | 0.95 | 94 |
| *daf-16* | *S. enterica* | 10 | 2.4e-2 |  | 0.13*** | -0.96*** | 1.42 | 40 |

Variation in infection was consistently high, and parametric susceptibility models did not perform particularly well when fitted to these data (**Figure 2**). The homogenous and heterogeneous models often provided similar fits to data (**Figure 2**) with similarly poor goodness of fit (**Table 2**), apparently due to the high variability in infection prevalence across exposure conditions. However, under homogeneous susceptibility, infection prevalence should approach 100% under the conditions used here, which ensured that all susceptible individuals were constantly exposed to the infectious agent. Worms feeding *ad libitum* in the highest bacterial densities should have experienced thousands of live, intact bacteria per hour entering the intestine (Vega and Gore 2017); as the capacity of the intestine is only ∼10^5^ bacteria, this is a heavy inoculum. Despite the heavy doses of bacteria and the long exposure times used, no condition produced 100% infections in any replicates; maximum infection prevalence observed in any population was 60%. Further, linear fits to low-exposure data (**Figure 2**, dotted lines; fit to data for exposure<100, time<=7, log_10_CFU<9) frequently under-or over-estimated infection prevalence at high exposures by a substantial margin. This suggests that, although both models provided poor fits to data, the heterogeneous susceptibility model should be preferred.

The combined exposure term *Pt* did not adequately represent the interaction between infectious dose and time in these data (**Figure 2A-B**), suggesting that exposure duration and dose are not interchangeable. Within each data set, the cumulative exposure term did not explain substantially more of the variation than each component of exposure (dose *P* or exposure duration *t*) individually. However, fits to bacterial density were in general worse, and the heterogeneous model frequently failed to converge with bacterial density as the independent variable, suggesting that duration of exposure was generally informative (**Table 2**).

The interaction between dose and exposure time differed between the commensal MYb14 and the pathogen *S. enterica*. For MYb14, infection prevalence was dose-dependent with short (2-7h) exposures and appeared to saturate with a long 24-hour exposure (**Figure 2, Figure S5**). In N2 hosts, prevalence of MYb14 infection depended on infectious dose, with infection prevalence increasing monotonically over 7-9 logs of bacterial density (all Bonferroni-corrected Dunns test comparisons p<0.05 in full data set and p<0.003 when 24 hour exposure is removed); this dose-dependence was weaker in *daf-16* hosts (corrected Dunn’s test comparison of 7 vs 9 log(CFU) p=0.003 with 24 hour exposure data removed; all other comparisons n.s.). Ignoring the effects of dose, accumulation of MYb14 infections was not distinguishable among short exposures (2-7h) but was distinct between long and short exposures in both hosts (Kruskal-Wallace p<4e-4, all corrected Dunn tests vs 24-hour exposure p<0.005, other comparisons n.s.).

In *S. enterica*, the relationship between infection prevalence and dose was dependent on exposure time (**Figure 2; Figure S5**). In both hosts, infections generally increased with exposure time (Dunn test comparisons of 3h vs 6-7h and 3h vs 24h all p<0.03; 6-7 vs 24 hours n.s.). Infection in N2 hosts was weakly dose-dependent overall (Kruskal-Wallace p=4.1e-4, comparison of 8 vs 10 log_10_CFU p=3.9e-4); this dependence was particularly marked at a long 24-hour exposure (Kruskal-Wallace p=8.9e-4, comparison of 8 vs 10 log_10_CFU p=3.97e-4) but was also present at short exposures (3-7 hour data only, Kruskal-Wallace p=0.022, comparison of 8 vs 10 log_10_CFU p=0.027, all other comparisons n.s). Infection in *daf-16* hosts was more strongly dose-dependent (Kruskal-Wallace p=1.4e-4, comparisons of 8 vs 9 and 8 vs 10 log_10_CFU p<0.03). In contrast to MYb14, long exposure times appeared to increase rather than decrease the effects of inoculum density in *S. enterica*. This is qualitatively consistent with expectations from a pathogen synergy model (Ben-Ami, Regoes, and Ebert 2008), where an increase in agent density can produce an over-proportionate increase in infections, but it is difficult to confirm this against the substantially sub-linear background of the overall dose-infectivity relationship.

Overall, these data indicated a complex relationship between infection and exposure which was affected by the properties of the infecting agent and the susceptible population. Variation in infection prevalence was consistently high across exposure doses and durations, and traditional models for susceptibility did not provide good fits to these data. However, the observed failure to approach 100% infection prevalence despite heavy inoculation, as well as the inability of the homogeneous model to predict infection prevalence outside of the range of the data, suggested that susceptibility to infection in *C. elegans* was not best described by a homogenous model.

### Secondary infections

By combining observed distributions of shedding and susceptibility, it should be possible to predict distributions of secondary infections. We therefore sought to describe distributions of secondary infections by *S. enterica* empirically, and then to predict these distributions using zero free parameter models for shedding and susceptibility from the previous sections. To obtain experimental distributions of secondary infections, we allowed N2 index cases (n=48) pre-infected with *S. enterica-GFP* to shed into liquid media, then transferred that supernatant onto susceptible populations for a pre-determined period of exposure (7h) to allow infection. After exposure, external bacteria were removed, and infections were allowed to develop as before (**Methods**) to allow detection via GFP fluorescence of individual worms. Supernatant from each index case was used to expose one susceptible population of each host genotype (N2, *daf-16*), allowing us to compare “copy-pasted” outbreaks across pairs of populations with different parameters for susceptibility. In this setup, standard assumptions for classical ODE compartment models (random mixing within compartments; stable external environment; constant host population size) are essentially true, allowing us to make direct comparisons between these data and the predictions of the simple models parameterized earlier.

To observe how different distributions of shedding and contact alter distributions of secondary infections, we investigated two transmission scenarios (**Figure 3**). Scenario 1 was an “everyone sheds” scenario where index-case supernatant was harvested at 24 hours, allowing most index cases to shed and resulting in a wide distribution of bacterial densities. Scenario 2 was a “super-shedding-biased” scenario where index-case supernatant was harvested at 7 hours, at which time most index cases had not shed bacteria. In the latter scenario, harvested index-case supernatant was incubated to 24 hours to allow bacterial growth, resulting in a bimodal distribution of infectious agent densities.

**Figure 3.**
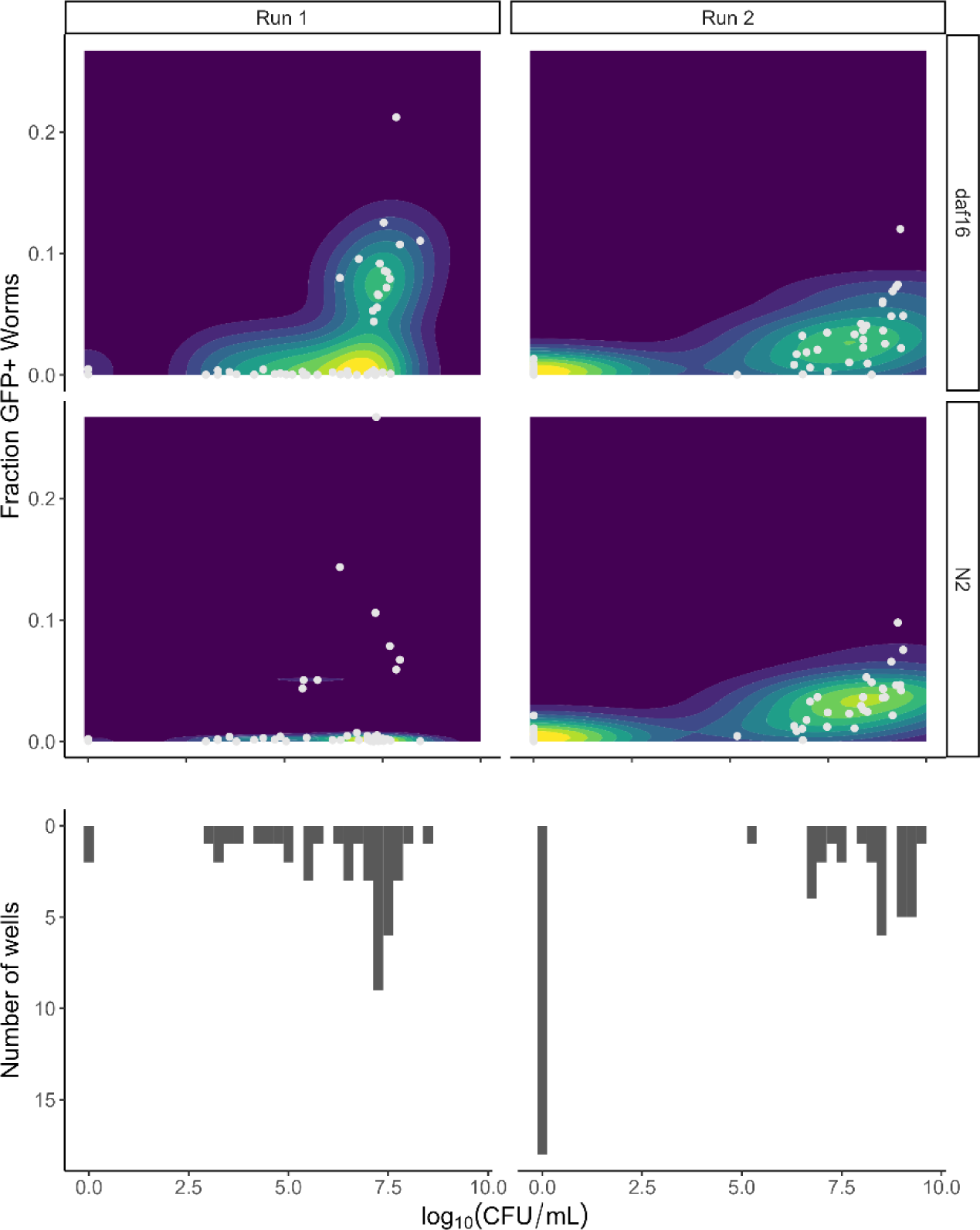
Secondary infections by *Salmonella enterica*-GFP from “copy-pasted” transmission events. (Top) Distributions of infection prevalence (fraction infected worms) vs density of the infectious agent. Contour color indicates relative density of outcomes (dark blue=low; yellow=high). Data points are shown in light grey. Each data point represents one replicate population in a single well. (Bottom) Histograms of bacterial densities to which susceptible worms were exposed in each scenario, measured by plating on solid agar for colony counts. Supernatant from 48 index case wells was harvested either (Run 1) after 24 hours shedding, to allow a full distribution of shedding events, or (Run 2) after 7 hours (+ out-growth to 24 hours), to restrict the duration of shedding. Supernatant from each index case was used to expose initially susceptible populations of N2 and *daf-16* worms (**Table 3**), so that each transmission event occurred in one pair of susceptible populations. Susceptible populations (run 1, n=15-20 susceptible worms/well; run 2, n=120-150) were exposed to the infectious agent for 7h, then moved to inert food to allow infections to develop. Infections were read on Biosorter 36 hours post-exposure, and individuals were scored as infected (GFP+) if they were above the 99^th^ percentile of green channel fluorescence obtained from non-fluorescent controls.

**Table 3.**
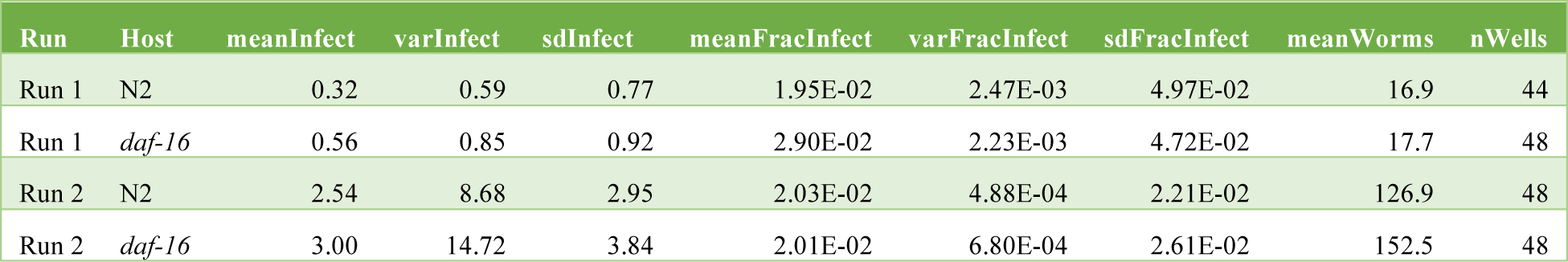
Summary statistics for secondary infections by *S. enterica*-GFP. *meanInfect* and *varInfect* are mean and variance of the number of infected worms (GFP+) across all wells in an experiment. *meanFracInfect* and *varFracInfect* are the indicated statistics for the fraction of infected individuals across wells. *meanWorms* is the average population size (total number of initially susceptible worms) across wells. *nWells* is the total number of populations measured in each experiment.

**Table 4.**
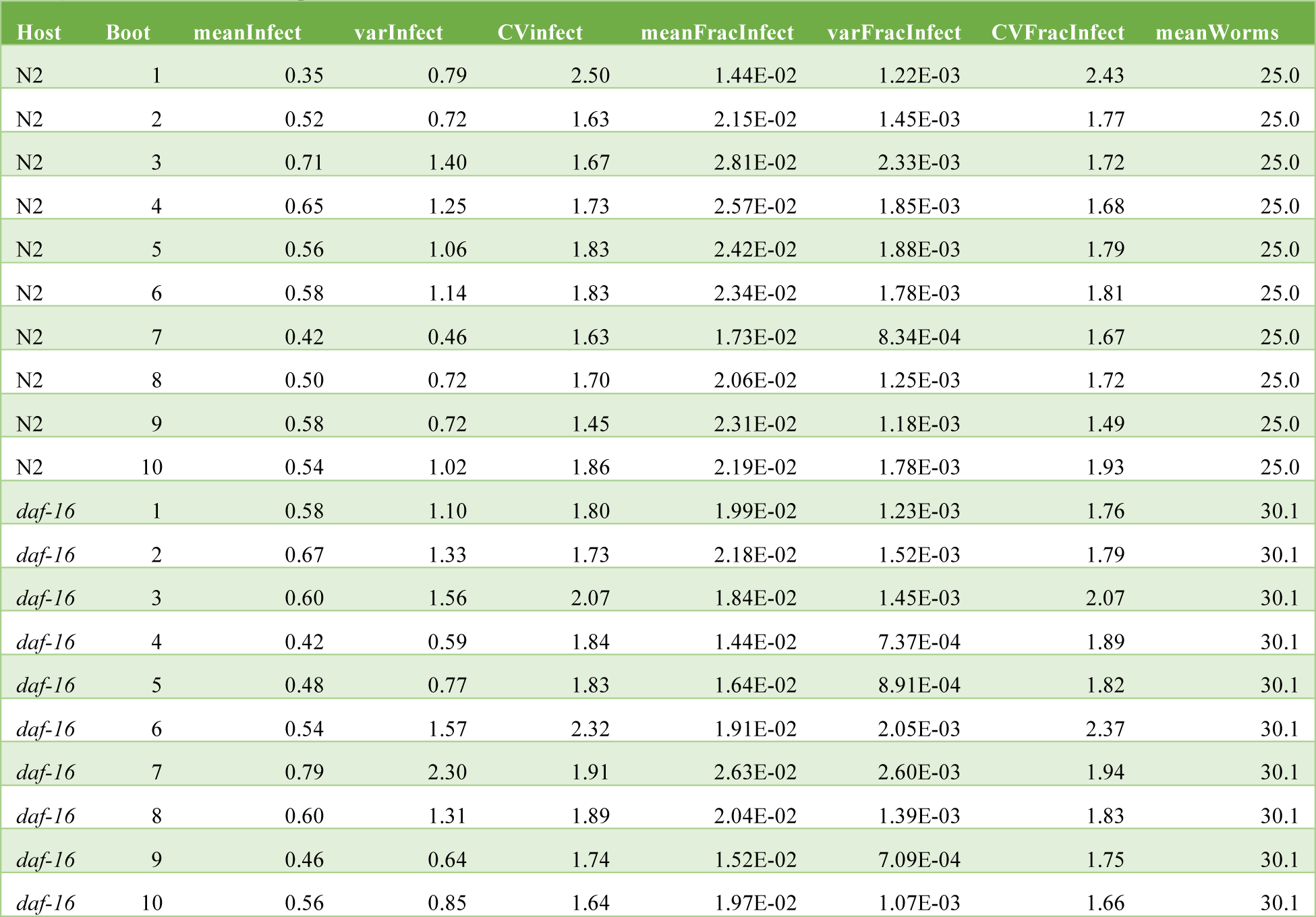
Summary statistics for resampled data, using 1/5 of the original worms per well. Bootstrapping was done on raw Biosorter data from run 2, with resampling within wells. *Boot* is the index of the bootstrap replicate (10 runs per host). *meanInfect*, *varInfect*, and *CVInfect* are statistics for the number of infected worms (GFP+) across all wells in the resample. *meanFracInfect*, *varFracInfect*, and *CVFracInfect* are the indicated statistics for the fraction of infected individuals across wells. *meanWorms* is the average population size (total number of initially susceptible worms) across wells in the resampled data.

As expected from previous results, infection prevalence increased with bacterial density and was highly variable across densities of the infectious agent (**Figure 3**). Most transmission events resulted in zero secondary infections, and the fraction of “outbreaks” with no secondary cases decreased as bacterial density increased. Number but not frequency of infections increased with susceptible population size (**Table 3, Figure S6**). Average frequency of infection was low; the “high-susceptibility” *daf-16* host had a higher maximum infection prevalence than N2, but the median for both hosts was 0 (**Table 3, Figure S6**). Variance in the observed frequency of infection was high (variance ∼ mean^2^ independent of susceptible population size, **Tables 3+4**).

We next attempted to predict these results using previously fitted models for *S. enterica* infections (**Figure 4**). First, we attempted to predict distributions of secondary infections based on known exposures, using measured bacterial densities from each well and the constant exposure duration (7 hours) used in these experiments. In these simulations, bacterial densities and number of susceptible worms for each population were copied from data (**Figure 3**). Secondary infections were simulated from the homogeneous (SI) and heterogeneous (Gamma) susceptibility models, using parameters for *S. enterica* infection in each host genotype (**Table 1**).

**Figure 4.**
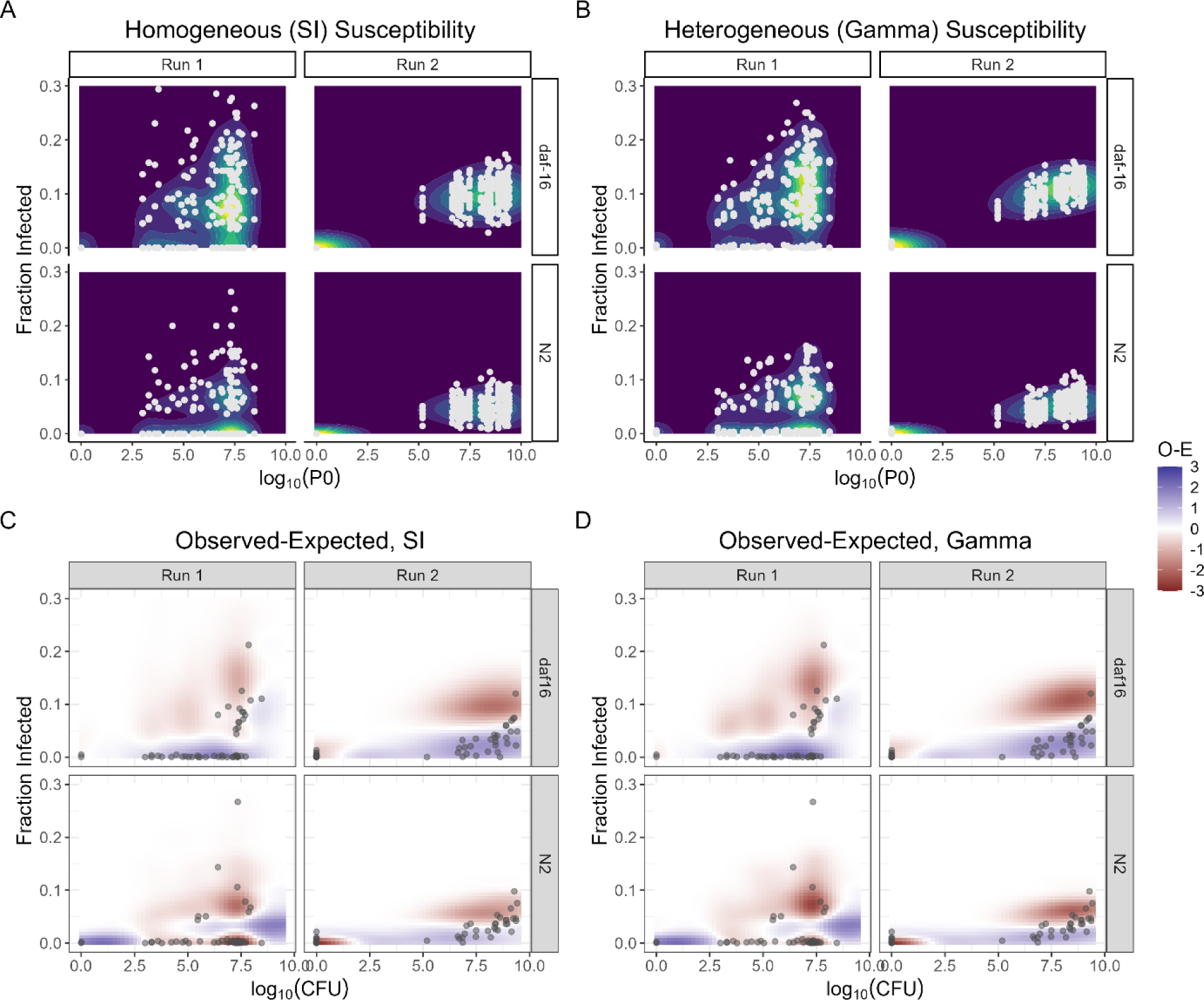
Model predictions for distributions of secondary infections by *S. enterica*-GFP, using bacterial densities and susceptible population sizes from data. Ten simulations were carried out at each combination of bacterial density and susceptible population size. (A-B) Simulation results for secondary infections with (A) homogeneous and (B) heterogeneous susceptibility models. Simulated data points are shown in light grey. Contour colors represent relative density of outcomes (low=dark blue, high=yellow). Simulation parameters for susceptibility models were taken from fits to *S. enterica* susceptibility data (Figure 2, **Table 2**); parameters for growth and uptake of bacteria were taken from shedding experiments (Figure 1) and from (Vega and Gore 2017). (C-D) Differences between observed and predicted density of outcomes, calculated as density of real data (observed) – density of simulation results (expected). Areas where experimental data had higher density of outcomes than simulations are in blue; areas where simulations predicted higher outcome density are in red. Experimental (observed) data points are shown in dark grey.

Model predictions in general over-estimated the frequency of infection (**Figure 4**). The SI model predicted excess infections at all bacterial concentrations used, including a prediction of non-zero infection rates at concentrations that were sub-infectious in the data (**Figure 4A,C**). The homogenous-susceptibility (SI) model fitted to only data for 6-7 hour exposure times performed considerably worse than the same susceptibility model fitted to the full data set (**Figure S7**). The heterogeneous susceptibility model performed somewhat better than the homogeneous model (**Figure 4B,D**), with fewer excess infections predicted, particularly at lower bacterial densities. Variation in fraction infected individuals decreased markedly in both sets of simulations when the number of susceptible hosts was increased (**Figure 4, Figure S8**), whereas the data indicated high variation in secondary infection rates even for large susceptible populations (run 2, **Figure 3**). Heavy inoculum doses (high bacterial densities) tended to decrease variation in infection prevalence in both experiments and simulations. Particularly for *daf-16*, simulation results tended to under-estimate the variation in infection at a given bacterial density (**Figure S6C**).

In these experiments, density of the infectious agent was measured prior to exposure and was available as a model input. However, in most cases exposure to the infectious agent cannot be measured directly and must be predicted. To obtain predictions for exposure intensity, we therefore simulated bacterial shedding using the homogeneous and Pareto models parameterized from earlier experiments, then fed the resulting predictions into simulations to predict secondary infections in N2 worms (**Figure 5**). For scenario 1, where index cases were allowed to shed for the full 24 hours, the constant-shedding model predicts substantially less spread, as well as a lower peak, in the distribution of bacterial densities as compared with the data; the Pareto-shedding model out-performed the constant-shedding model on both counts (**Figure 5A**). For scenario 2, where index case shedding was stopped after seven hours and bacteria were then allowed to grow in the absence of new shedding, both models predict similar distributions of agent density; the waiting-time distribution for shedding is conserved across models, and with this scenario, the number of index cases that have successfully shed before the stopping time is largely responsible for the distribution of outcomes. The effects on predicted distributions of secondary infections were as expected, with very little difference between shedding models. As before, heterogeneous (Gamma) susceptibility provided the best matches to data, particularly at lower densities of the infectious agent, but differences between models were small (**Figure 5B**).

**Figure 5.**
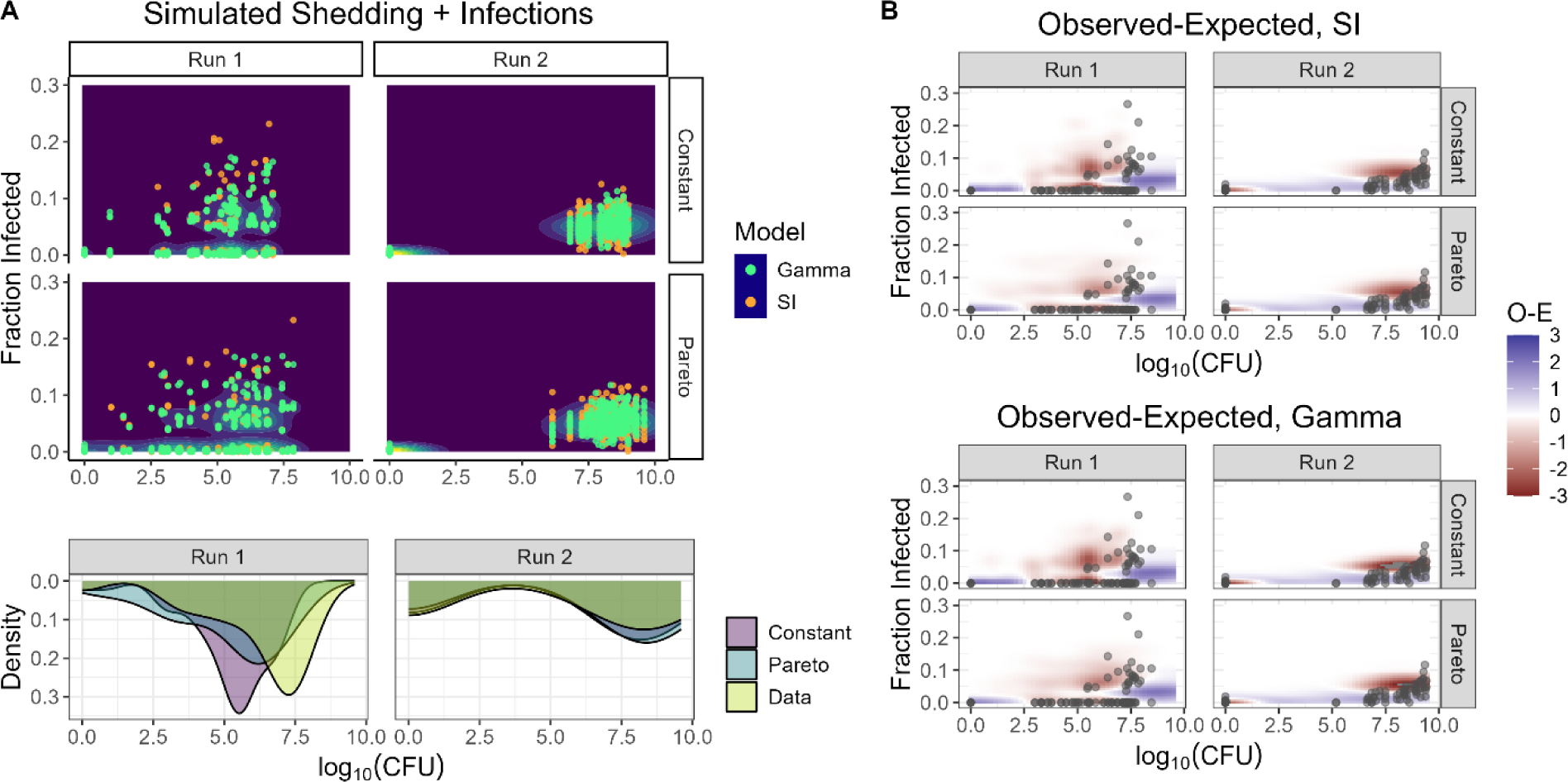
Model predictions for distributions of secondary infections in N2 worms by *S. enterica*-GFP, using susceptible host population sizes from data and simulating bacterial densities from shedding models (Poisson rate; constant or Pareto magnitude). Ten simulations were carried out at each combination of bacterial density and susceptible population size. Simulation parameters for susceptibility models were taken from fits to *S. enterica* susceptibility data (Figure 2, **Table 2**); parameters for growth and uptake of bacteria were taken from shedding experiments (Figure 1) and from (Vega and Gore 2017). (A) Simulated distributions of secondary infections (top) resulting from simulated bacterial densities after shedding (bottom; bacterial densities observed in experiments shown as “Data” in green for comparison). (B) Differences between observed and predicted density of outcomes, calculated as density of real data (observed) – density of simulation results (expected). Results from the homogeneous (SI, top) and heterogeneous (Gamma, bottom) susceptibility models are shown separately. Areas where experimental data had higher density of outcomes than simulations are in blue; areas where simulations predicted higher outcome density are in red. Experimental (observed) data points are shown in dark grey.

## Discussion

As in other systems, microbial transmission in the nematode *Caenorhabditis elegans* is highly variable. We observed that shedding rates for gut-associated bacteria in this host are low, consistent with previous work (Taylor, Spandana Boddu, and Vega 2022), and that these low rates result in wide waiting-time distributions for shedding events. Super-shedding was common, occurring at frequencies similar to those inferred for infectious diseases in other hosts. Accumulation of new infections in this host was likewise variable, with exposure time and exposure intensity explaining some of this variation, but with considerable residual variation across replicate populations exposed to identical conditions in parallel. When susceptible populations were exposed to shedding from index cases, most transmission events resulted in no secondary infections as expected. Further, distributions of secondary infections were poorly predicted when homogeneous shedding and/or susceptibility was assumed, again consistent with inferences from real-world transmission networks. Our results suggest that transmission in *C. elegans* shares conserved features with transmission in more complex systems.

Our observations support the idea of super-shedding as a biologically intrinsic property of transmission. This system represents possibly the simplest, most uniform possible scenario, where highly homozygous, age-synchronized index cases arising from a single, shared exposure to a clonal agent are allowed to shed in parallel in a highly controlled environment, and where shedding is measured directly rather than via secondary infections. Even in this stripped-down scenario, we observed super-shedding by a consistent and familiar fraction of index cases. This is not to say that other aspects of super-shedding, in particular behavior within transmission networks, are not important. Even in this simplified system, where the only conventional “behavior” permitted is the (operator-controlled) number of susceptibles per exposure, this “behavior” clearly matters for the number of secondary infections. Instead, our results support the idea that Pareto-type super-shedding should be considered when describing biological variation in shedding.

We observed substantial variation in accumulation of new infections when replicate populations were given the same or very similar exposures to an infectious agent. Observed variation was often in excess of stochastic model predictions (demographic noise), in models with and without population heterogeneity. Given the highly controlled nature of these experiments, this suggests that some source(s) of noise and/or heterogeneity in transmission are not accounted for. This is despite the fact that the explicit and implicit assumptions of canonical compartment models (well-mixed, essentially time-invariant, all-meets-all environments; constant population sizes; highly uniform cohorts of hosts) are very nearly true in these experiments. Some of this variation may be attributable to heterogeneity among hosts that is not captured by the models. Even in age-synchronized populations of worms from a single genetic background, individuals differ in stress tolerance, lifespan, and rates and durations of feeding (Taylor, Spandana Boddu, and Vega 2022; Brooks, Lithgow, and Johnson 1994; Chen, Zajitschek, and Maklakov 2013; Natesan et al. 2023; Wu et al. 2006; Suda, Shoyama, and Shimizu 2009; Lee et al. 2017); some of this variation is due to differences in life history (Perez et al. 2017; Willis, Sukhdeo, and Reinke 2021), but genetic, epigenetic, and developmental sources of variation may also contribute (Rockman, Skrovanek, and Kruglyak 2010; Perez et al. 2017; Jobson et al. 2015). The relative contributions of different aspects of individual heterogeneity to population-level variation in transmission outcomes remain to be determined.

Canonical models for disease dynamics fail on these data in familiar ways, and ability to choose the best-performing model *a priori* was limited by the properties of the data used to make this decision. The homogeneous (SI) model for susceptibility over-estimated accumulation of secondary infections particularly at low exposures (**Figure 4**); this is well known to occur when a homogeneous model is used to make predictions for a heterogeneous population (Lloyd-Smith et al. 2005). This was true even though the two susceptibility models could not generally be distinguished based on goodness of fit in independent susceptibility experiments, in part due to the high variation in these data (**Figure 2**). The homogeneous susceptibility model performed particularly badly outside the range of the fitted data, as is often the case for regressions; further, a homogeneous model fitted to very limited data had little predictive value even though the scenario being predicted was within the range of the fitted data (**Figure S7**). Likewise, a constant-shedding model could be out-performed by one that allowed Pareto events, but the difference between models was small when shedding was externally constrained such that timing was more important than magnitude (**Figure 5**). In this scenario, where susceptible populations either had no exposure or were exposed to high densities of the agent for a fixed duration, but where infection prevalence was low (mostly <10%), differences between susceptibility models were also predictably small. The observation that data limitations can affect model choice is not a new one; our results are an illustration of this general principle, and the specific effects we have observed may be of broader interest.

Our zero free parameter models in general over-estimated secondary infections (**Figures 4-5**). This might suggest run to run differences in average host susceptibility. However, we observed a similar deviation between observed and expected results in two runs of this experiment conducted over a year apart, and it is improbable that we would obtain the same batch effect in both experiments. It is more likely that this observation reflects a difference in infectivity when *S. enterica* is cultivated *in vitro* (**Figure 2**) vs shed from index cases (**Figure 3**). Growth temperature was held constant at 25°C, but differences in the growth medium (NGM vs S-medium + heat-killed OP50) could result in phenotypic differences, as could passage through an index case host. This is a caveat as well as a potential opportunity for future work.

These experiments were intended as a proof of concept for the *C. elegans* model, where the assumptions of traditional compartment models were met as nearly as possible to allow direct comparison between experimental observations and the predictions of those models. As such, certain important features of real-world epidemics were excluded. This was a deliberate choice to maintain simplicity, and features can be added to these experiments as desired. For example, while we began with pure inbred host lineages and clonal bacterial stocks, addition of genetic diversity to the starting conditions could be used to examine the effects of variation in host immunity and/or agent traits on outcomes during transmission.

Further, behavior is a critical component of transmission in real epidemics which was deliberately not allowed to vary in these experiments. The simplest relevant component of behavior is connectivity – the number of susceptible individuals encountered by a given index case while infectious – which may or may not be correlated with susceptibility in a given scenario (White, Forester, and Craft 2018). With the current experimental setup, where shedding and transmission are separated, it would be straightforward to introduce this “behavior” by altering the number of susceptibles for each transmission event according to the desired correlation(s). Further, exposure intensity in the real world can be separated into frequency and duration of contacts, which are properties of the social network (Bansal, Grenfell, and Meyers 2007); this could likewise be varied experimentally, but for simplicity in the proof of concept, we chose to keep susceptible hosts in continuous contact with the agent and vary only the duration of exposure. Finally, it is not at present possible to determine whether an index case will be a super-shedder *a priori*, without measuring shedding after the fact. Nor is it actually clear that “super-shedding” is a property of the individual, as the effects of shedding events after the first successful shed will be swamped by exponential growth in these assays. It is possible that all individuals have a probability of super-shedding, such that the Pareto distribution for shedding is sampled by individuals across shedding events rather than by the population of index cases across individuals. These ideas remain as challenges for future work.

The experimental approaches used here were designed to quantify transmission of fluorescently labeled bacteria in *C. elegans* but could easily be generalized. Use of agents labeled with a fluorescent reporter allowed high-throughput measurement of intestinal bacterial load on a large object sorter, which is convenient but not strictly necessary; quantification of bacterial load in individual worms via destructive sampling (Taylor, Spandana Boddu, and Vega 2022) requires more effort per data point but can be used to measure unlabeled agents. Additionally, host reporters could be used alongside or instead of labels on the infectious agent if host response (e.g. immune induction) is of primary interest, or if this is easier to observe than microbial load. The protocol used here to measure shedding from supernatant is useful for agents that grow logistically in media with a negligible lag time post-shedding; this is reasonably common but is not true for all bacteria, as shown here by the example of *S. aureus*, where an extended lag phase prevented calculation of waiting times and shedding rates. For agents that do not show smooth logistic growth, or which do not grow outside the host, a qPCR-based approach might be preferable.

In these experiments, we investigated shedding and transmission of gut-colonizing bacteria that grow in an environmental reservoir before transmission to new hosts. Many of the agents commonly used in *C. elegans* host-microbe association fit this description, and an improved understanding of within-and between-host dynamics may introduce new dimensions to studies of these agents. For example, whereas response of *C. elegans* to an agent is generally measured as physiological response (e.g. lifespan, healthspan, fecundity) during continuous exposure to heroic doses of the agent (Park, Jung, and Lee 2017), there is increasing interest in understanding host response in more naturalistic scenarios where exposure to the agent is not constant. Further, there is interest in understanding how nematode and bacterial populations interact in structured environments such as their native soil habitat. These experiments used a well-mixed liquid environment to measure baseline parameters, which can be combined with spatial information (e.g. chemotaxis, decision-making, aggregation, “farming” on surfaces) (Thutupalli et al. 2017; Lee et al. 2017; Duckett et al. 2024; Demir et al. 2020) to gain a more complete understanding of how worm and bacterial populations interact in structured environments.

Other agents, particularly intracellular pathogens like microsporidia and the Orsay virus of *C. elegans*, may represent a quantitatively distinct transmission scenario (Frézal et al. 2019; Vassallo et al. 2023; González and Félix 2024; Castiglioni et al. 2024; Shaw and Kennedy 2022). Among other factors, these agents decay rather than proliferating in the environment. Quantifying shedding and transmission of these agents is more challenging than for bacteria, as plating-based live-count assays are not possible. Nonetheless, considering shedding and transmission of these agents as stochastic processes may reveal interesting differences in the distributions of transmission by agents with different modes of infection.

The current work used a simplified system where the demographic complications (e.g. overlapping generations, non-constant populations, genetically heterogeneous host populations), spatial considerations, and evolutionary dynamics of real-world infections were explicitly disallowed. From this point, complications can be added as needed to address specific questions. The work shown here is intended as a starting point for use of *C. elegans* as a tractable and quantitative transmission system.

## Acknowledgements

The authors would like to acknowledge the Civitello, Koelle, and Moran labs at Emory University for productive discussions and feedback on this manuscript.

## AI Usage

The author used ResearchRabbit.ai to accelerate literature discovery. No AI tools were used during the preparation of the manuscript.

## Data Availability

Data and code for this manuscript are archived on Emory Dataverse (https://doi.org/10.15139/S3/6J8JTI).

## Methods

### Strains and cultivation

Bacterial strains used in these experiments were *Salmonella enterica* LT2 with GFP and a kanamycin resistance cassette chromosomally integrated at the neutral *attB* site (Vega et al. 2013), *Staphylococcus aureus* Newman expressing GFP from a low-copy plasmid (pTRKH3-ermGFP, Addgene #27169) (Lizier et al. 2010), and *Ochrobactrum* MYb14 expressing GFP from a high-copy broad host range plasmid (pBTK519, KmR, Addgene #110603) (Leonard et al. 2018). *Escherichia coli* OP50 (CGC) was used as a standard food source; heat-killed OP50 was used as an inert food.

*Caenorhabditis elegans* strains were provided by the CGC, which is funded by NIH Office of Research Infrastructure Programs (P40 OD010440). Strains used in these experiments were wild type N2 Bristol and *daf-16(mu86)* I (CGC strain CF1038). Unless otherwise stated, worms were cultivated on 10 cm NGM agar plates at 25°C with *E. coli* OP50 as a food source using standard protocols (Stiernagle 2006).

### Bacterial shedding

Synchronized, reproductively sterile adult N2 worms were prepared according to standard protocols for this lab (Taylor, Spandana Boddu, and Vega 2022) and colonized for 16-48 hours on 10 cm nematode growth agar (NGM) plates with pre-established bacterial lawns covering the entire surface of the plate. After colonization, worms were rinsed thoroughly in M9 worm buffer + 0.1% Triton X-100 (M9TX01) to remove the bulk of remaining bacteria, incubated in S medium + 2X heat-killed *E. coli* OP50 for at least one hour to purge non-adhered bacteria from the gut, and run through a large object sorter (Biosorter, Union Biometrica, 250FOCA) to separate worms into bins based on GFP fluorescence from intestinal bacteria (**Figure S1**). Non-fluorescent worms (OP50 only) were used to determine thresholds for detectable colonization, such that the bottom of the “low” fluorescence bin was defined by the 95^th^ percentile of auto-fluorescence. Worms were then chilled to halt feeding and excretion, lightly surface bleached (1:2000 bleach in cold M9TX01 buffer for 20 minutes) to kill external bacteria, and rinsed 3X in cold M9TX01 to remove bleach. Worms were manually pipetted while still cold-paralyzed into individual wells of a 96-well plate (n=16-24 worms per fluorescence bin) containing 200 µL worm S medium + heat-killed OP50 as an inert food source + 10 µg/mL nystatin (to prevent fungal contamination) + selection appropriate for the colonizing bacterial strain (*S. enterica*, 25 µg/mL kanamycin; *S. aureus*, erythromycin 10 µg/mL; MYb14, 50 µg/mL kanamycin). Index case plates were covered with a BreatheEasy gas-permeable membrane and incubated at 25°C with shaking at 200 RPM.

Excretion of live bacteria was monitored by plating 5 µL volumes of undiluted supernatant at each time point, generating count data from which the time of the first shedding event could be back-calculated. Briefly, we calculated time since N_0_=3 (*WaitTime*) from the first sample where bacterial density became countable in each well (>4 colonies), assuming that bacterial growth at these low densities is exponential. (Carrying out the same calculation with N_0_=1 provided similar results, but the estimator was more stable with the more lenient initial condition, particularly for runs where the exponential growth rate of the bacteria was high.) We neglected additional shedding events under the assumption that their contribution is much smaller than that of growth when the shedding rate is small. We then calculated the rate parameter λ for a Poisson process as *1/mean(WaitTime)*, assuming that the inferred times of first shedding represent an exponential waiting time process. The calculated standard deviation of these waiting times was similar to the inferred SD for the resulting exponential, easily attributable to measurement error. In some wells, bacterial density increased too quickly to be the result of exponential growth from a single live bacterium in the elapsed interval; these worms were considered “super-shedders” and were not used for waiting time calculations. Control wells (n=3) containing buffer from which the infected worms were taken remained sterile, indicating that bacterial growth was due to shedding from live worms.

After 24 hours, worms were retrieved from individual wells by pipetting under a dissecting microscope. Wells for which no intact, live worm was found were removed from analysis to avoid treating wells that had not received a worm, or where the worm had died, as data. In a subset of experiments, bacterial load was confirmed in retrieved worms by disrupting these worms using standard protocols (Taylor, Spandana Boddu, and Vega 2022) and plating whole-gut contents for CFUs.

Gillespie simulations for time to detectability in index worm supernatant were carried out in R using *GillespieSSA::ssa()* with the tau-leaping method. Parameters for bacterial growth were inferred from data within each run of the experiment; death rate of bacteria in the supernatant was assumed to be low (0.05 h^-1^). For simulations of time to detectability during shedding in individual wells, carrying capacity K was held at 10^4^ bacteria to minimize run time; as detectability occurs far below this threshold, this value is arbitrary.

Jackknife confidence intervals for the rate parameter and for the fraction of super-shedders were calculated from data within each run using *bootstrap:: jackknife()*. Pareto fits to fraction detectable wells were carried out using *EnvStats::optim()* to minimize SSE of data vs. Gillespie simulation output, with number of bacteria in a shedding event distributed as Pareto with location = 3 and fitting the shape parameter α.

### Susceptibility vs known bacterial dose and exposure duration

Germ-free, synchronized, reproductively sterile populations of adult worms were prepared as previously described. These initially susceptible populations were resuspended in S medium + antibiotic (25 µg/mL kanamycin for selection on *S. enterica-*GFP, or 50 µg/mL kanamycin for MYb14-GFP) + 10 µg/mL nystatin (to prevent fungal contamination), then distributed in 100 µL aliquots among wells of a 1.2 mL deep well 96-well plate (1-6 replicate wells per experiment, 30-100 worms/well).

Bacteria were grown to stationary phase in 2 x 2 mL LB + antibiotic for selection, then centrifuged 2 min @ 9000xg to pellet cells, washed once in 2×1 mL S medium, centrifuged again, and resuspended in 2×1 mL S medium + antibiotic for selection. This 2X suspension was diluted further into the same medium to achieve the range of bacterial densities required for a given experiment. To each well containing susceptible worms, 100 µL of the indicated suspension of fluorescently labeled bacteria was added, to achieve final bacterial concentrations in the range 10^7^-10^10^ CFU/mL. Control wells were given 2X HKOP50 in the same medium instead of live bacteria. Plates were covered with gas-permeable BreatheEasy membrane and incubated at 25°C with shaking at 200 RPM.

After an initial exposure of pre-determined duration (2-24 hours), the bulk of the supernatant was removed by pipetting. Worms were washed 2X in cold M9TX01, lightly surface-bleached as previously described to remove any remaining external bacteria, then rinsed 2X in M9TX01 and 1X in S medium to remove bleach. Worm populations from each well were then moved to the same well of a clean 96-well plate in 200 µL S medium + 2X HKOP50 + selective antibiotic + 25-50 µg/mL chloramphenicol to prevent re-inoculation during out-growth of the initial infection. Plates were covered with a gas-permeable BreatheEasy membrane and incubated at 25°C with shaking at 200 RPM. After 24-48 hours of out-growth, worms were washed 3X in M9TX01 to remove heat-killed bacteria, then run on the large object sorter (BioSorter) to determine GFP channel fluorescence in individual worms. Within each run, a threshold for infection was set at the 95^th^ percentile of fluorescence in un-infected controls (no exposure to fluorescent bacteria).

Homogeneous and heterogeneous susceptibility models were fitted to data, using bacterial density (log_10_(CFU)), exposure duration (in hours), or a combined exposure term (density * duration) as the independent variable and −*ln*(*S*(*t*)⁄*S*_0_) as the dependent variable as in (Dwyer, Elkinton, and Buonaccorsi 1997). The homogeneous SI model w gives a linear relationship between log(fraction infected) and time, −*ln*(*S*(*t*)⁄*S*(0)) = β*Pt*, where S(0) and S(t) are uninfected (susceptible) populations at time 0 and at time *t*, *β* is a transmission constant, and *P* represents intensity of exposure to the infectious agent (e.g. encounters with infected individuals or contaminant density in the environment); fits were carried out in R using function *lm()*. In the heterogeneous model used here, *β* is a Gamma r.v. with dispersion *k*, and the resulting function 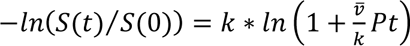 was fitted using *nls*() with starting parameters k=1.2 and 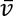=0.02. When *nls*() failed to converge, parameters for the Gamma distribution were estimated using *EnvStats::optim()* to minimize SSE between observed and predicted values for the dependent variable with method L-BFGS-B and parameter bounds 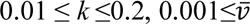 ≤0.02.

### Secondary Infections

N2 index cases and populations of susceptible hosts were prepared as previously described. Index cases were selected and sorted out based on membership in a wide “GFP+” gate (lower bound > upper bound of green channel for non-fluorescent control worms), then surface bleached and manually distributed among wells of a 96-well plate as previously described. In scenario 1 (“everyone sheds”), index cases were permitted to shed into 200 µL/well S medium + 1X heat-killed OP50 + selection (25 µg/mL kanamycin) + 10 µg/mL nystatin (to prevent fungal contamination) for 24 hours, after which the bulk of the supernatant was removed and wells were checked to confirm the presence of live, intact index case worms. 48 wells were chosen at random from the list of wells containing confirmed live index cases, and 50 µL of supernatant from each of these wells was added to wells of a deep 1.2 mL 96-well plate containing a population of susceptible hosts (one well each N2 and *daf-16*) in 150 µL S medium + 1X heat-killed OP50 + selection. An aliquot of the remaining supernatant was serially diluted and plated onto nutrient agar to determine bacterial density in each index case well. In scenario 2, index cases were set up as described, and a 5 µL aliquot of supernatant was taken at 7h and used to inoculate a second plate containing 195 µL/well of the same medium; this second plate was also covered with a BreatheEasy gas-permeable membrane and incubated at 25°C with shaking at 200 RPM, then used as an inoculum source at 24 hours as described for scenario 1.

In both scenarios, the 96-well plate of susceptible populations was covered with a BreatheEasy gas-permeable membrane and incubated at 25°C with shaking at 200 RPM for seven hours to allow infection to proceed. Non-fluorescent controls (heat-killed OP50 only) were maintained in a separate 96-well plate to prevent cross-contamination. After the infection period, the bulk of the supernatant was removed, wells were washed 2X with 1 mL M9TX01 to remove the bulk of the bacteria, and populations were chilled and surface-bleached to remove any remaining external bacteria. After bleaching, populations were rinsed 2X with 1 mL M9TX01 and 1X with 1 mL S medium, then resuspended in 200 µL S medium + 1X heat-killed OP50 + selection + 25 µg/mL chloramphenicol to prevent proliferation of *S. enterica* outside the host. Infections were read on Biosorter at 36 hours as GFP fluorescence in individual worms.

Gillespie simulations for bacterial shedding and secondary infections were carried out in R using *GillespieSSA::ssa()* with the tau-leaping method. Parameters for bacterial growth (exponential growth rate, carrying capacity) were inferred from data within each run of the experiment; death rate of bacteria in the supernatant was assumed to be low (0.05 h^-1^). Parameters for shedding (rate, Pareto α) and susceptibility (β or {k, v}) were taken from model fits in previous sections.

