## Supplemental File S1 for "Heterogeneous shedding and susceptibility in a *Caenorhabditis elegans* transmission model"

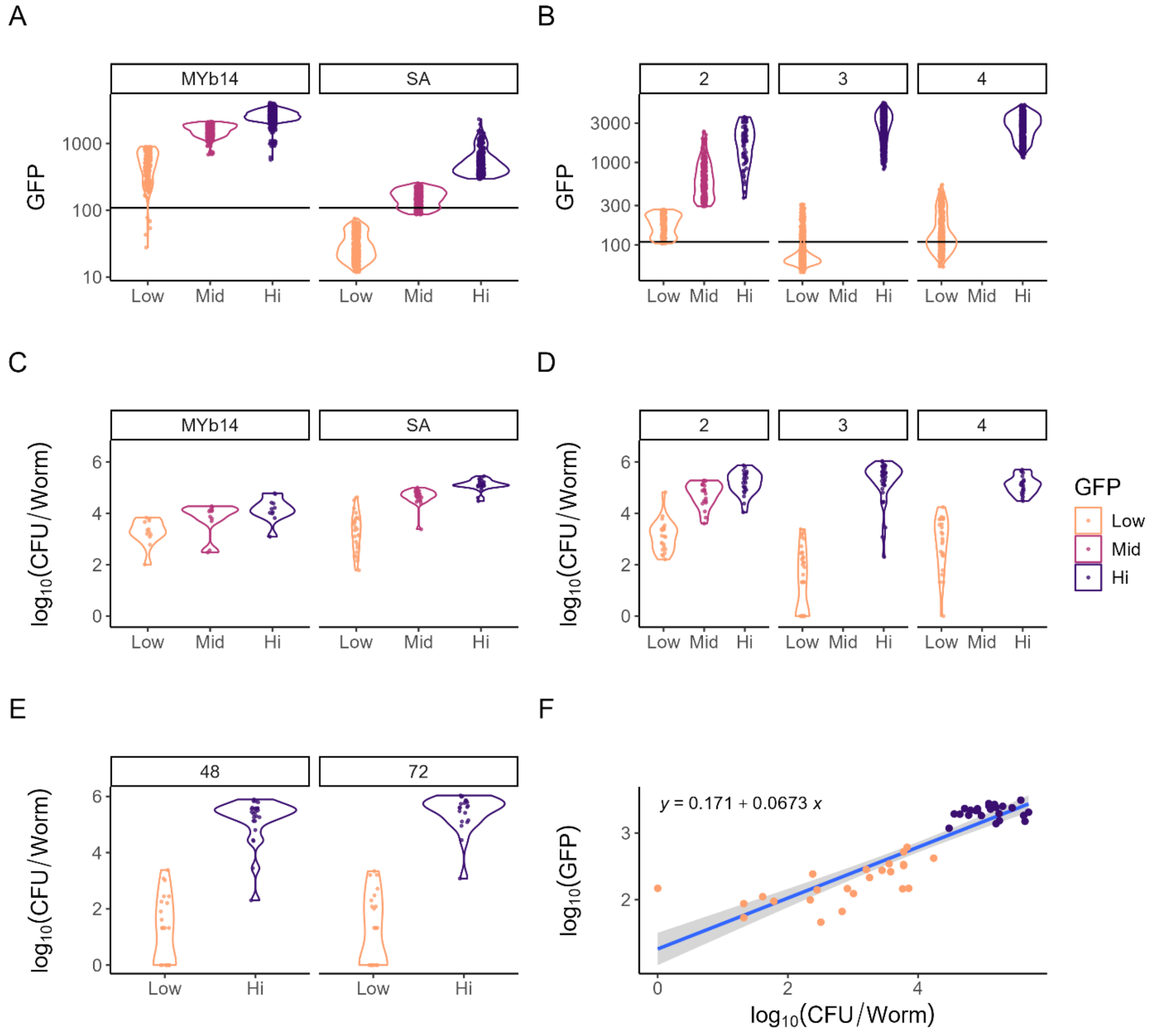


Figure S1. Bacterial fluorescence is a reliable proxy for bacterial load in individual worms. (A-B) GFP fluorescence and (C-D) bacterial load in individual worms ($\log_{10}$(CFU/worm)) within GFP bins, for (A, C) *Salmonella enterica*-GFP runs 2-4 and (B, D) other GFP-labeled bacterial agents *Ochrobactrum* MYb14 and *Staphylococcus aureus* Newman. Run 1 with *S. enterica* did not have CFU data and is omitted from these plots. Worms were run on BioSorter at the start of the experiment to separate individual worms into bins based on GFP fluorescence (low/mid/high); a sub-sample from each bin (n=12-36 worms) was reserved for single-worm disruption and plating to determine CFU/worm. Black line represents 90$^{th}$ percentile GFP fluorescence of control worms (given unlabeled *E. coli* OP50) from the same experiment. Note that in some runs 3-4 the “low” gate was set up to include some individuals below the apparent threshold of detection. These data suggest a GFP threshold of detection of ~100 bacteria/worm for *S. enterica*-GFP (integrated cassette), ~100 CFU/worm for MYb14-GFP (high-copy plasmid) and ~$10^4$ CFU/worm for *S. aureus*-GFP (low-copy plasmid). Plating TOD is ~40 bacteria/worm for all experiments (200 µL volume, plating 10 µL spots). (E) Duration of exposure to *S. enterica*-GFP does not alter distributions of bacterial load per worm. (F) CFU to GFP mapping for individual worms colonized with *S. enterica*-GFP. To obtain these data, individual worms were sorted directly into wells for single-worm disruption and plating, such that Biosorter measurement of total GFP fluorescence and CFU data for individual worms had the same well ID. In all plots, each point represents a single worm.

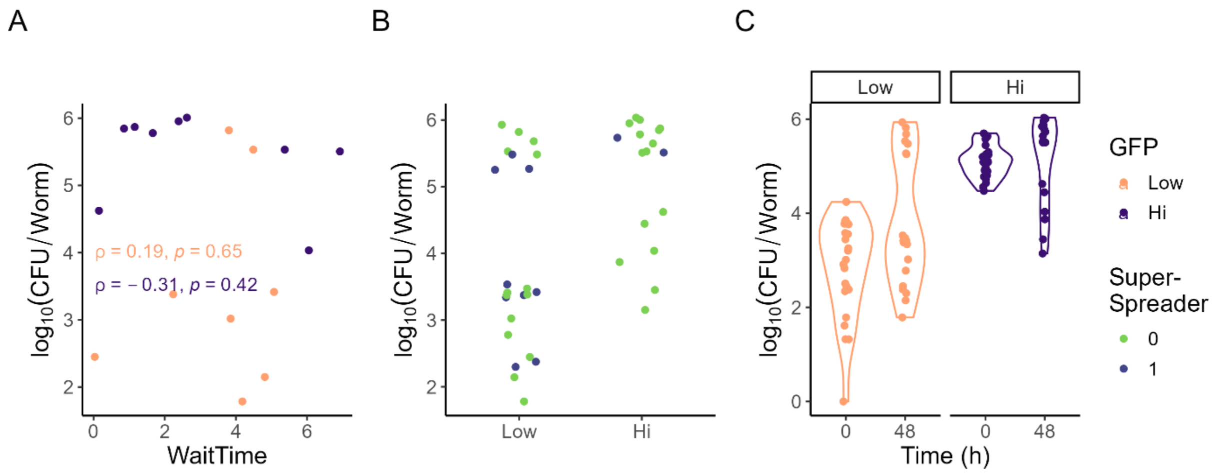


Figure S2. Bacterial fluorescence of *S. enterica*-GFP in individual worms does not predict shedding and can change over time. Each point represents one individual worm from run 4. (A) *WaitTime* is not significantly correlated with bacterial load in individual worms. CFU/worm was measured by disruption and plating of worms retrieved from wells at the end of the shedding experiment; each data point represents a well for which a live, intact worm was retrieved and for which *WaitTime* could be estimated. (B) Super-spreader classification of worms by initial GFP bin and bacterial load at the end of the experiment. Note that although worms were sorted into the GFP bins at time 0, some worms moved between bins in total bacterial load over the intervening 48 hours; this is typical. (C) Bacterial load can change over time in worms colonized by *S. enterica*-GFP, as can be seen by comparing bacterial load from worms sorted into each fluorescence bin at time 0 (experiment setup) with that of worms retrieved at the end of the experiment (48 hours). Note that sampling for CFU quantification is destructive; time-0 and time-48 worms are different individuals from the same populations. Data are colored by (A, C) GFP bin or (B) super-shedder status.

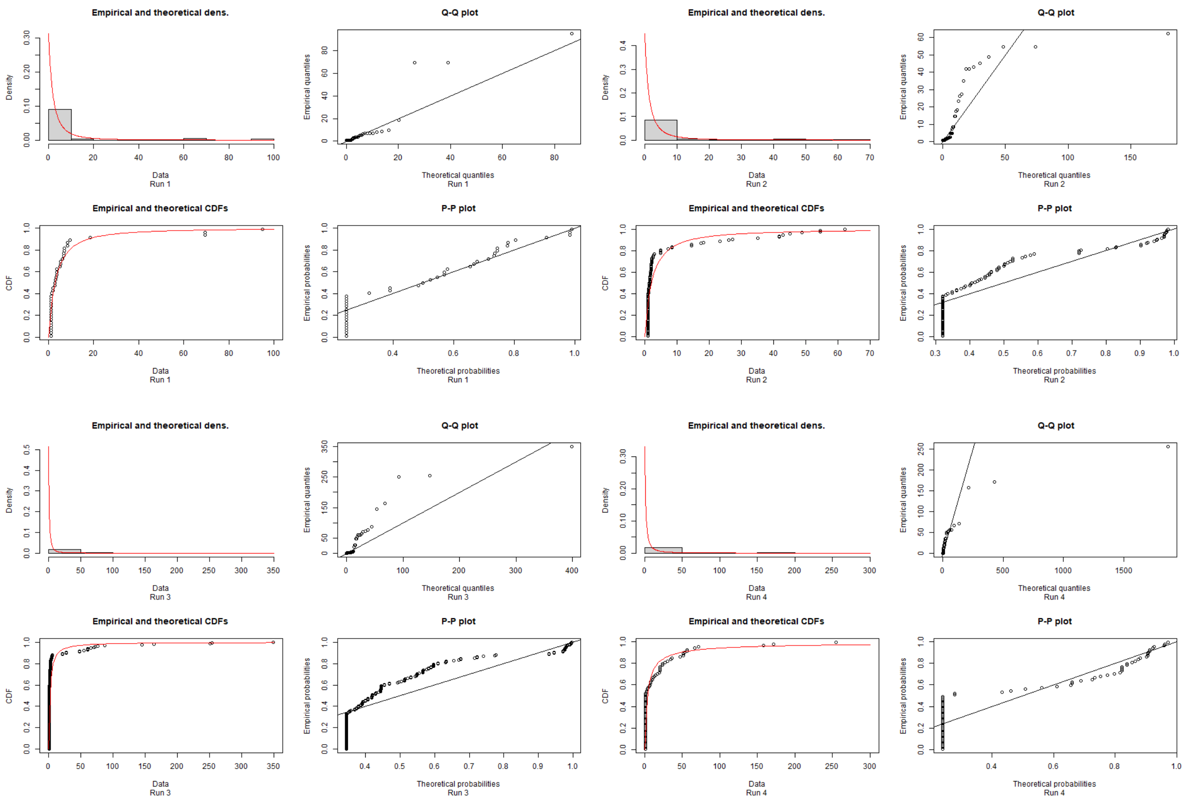


Figure S3. Pareto fits to inferred number of bacteria per shedding event for *S. enterica*. Data from each run (1-4) were fitted separately using function *fitdist* from R package *fitdistrplus*; Pareto distribution is provided in package *actuar*. Fitting was initiated with parameters shape=1, scale=2.

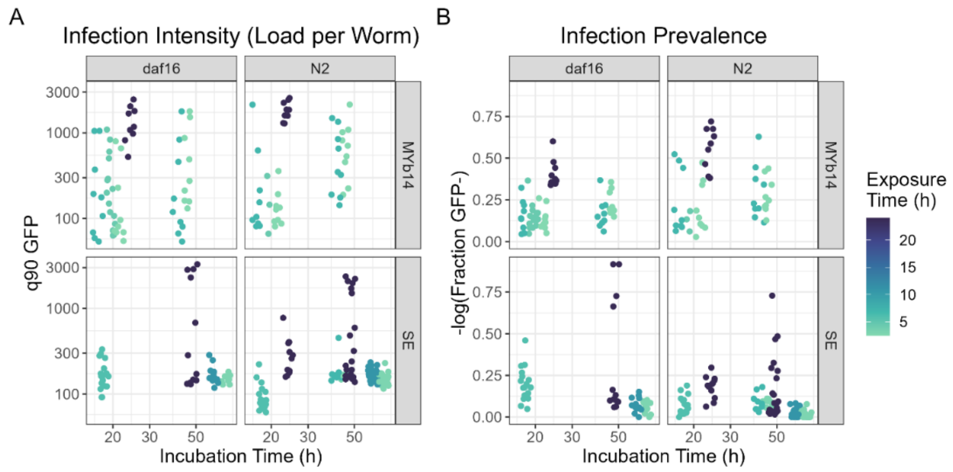


Figure S4. Accumulation of new infections is largely unaffected by duration of out-growth (incubation time) post-exposure. Data shown are incubation time in hours vs (A) peak infection intensity (90$^{th}$ percentile of GFP fluorescence across worms within a sample) and (B) infection prevalence within samples. Each point is one replicate well (30-100 worms) from the data set in **Figure 2**. Colors indicate duration of exposure to the infectious agent prior to incubation on inert food + antibiotics.

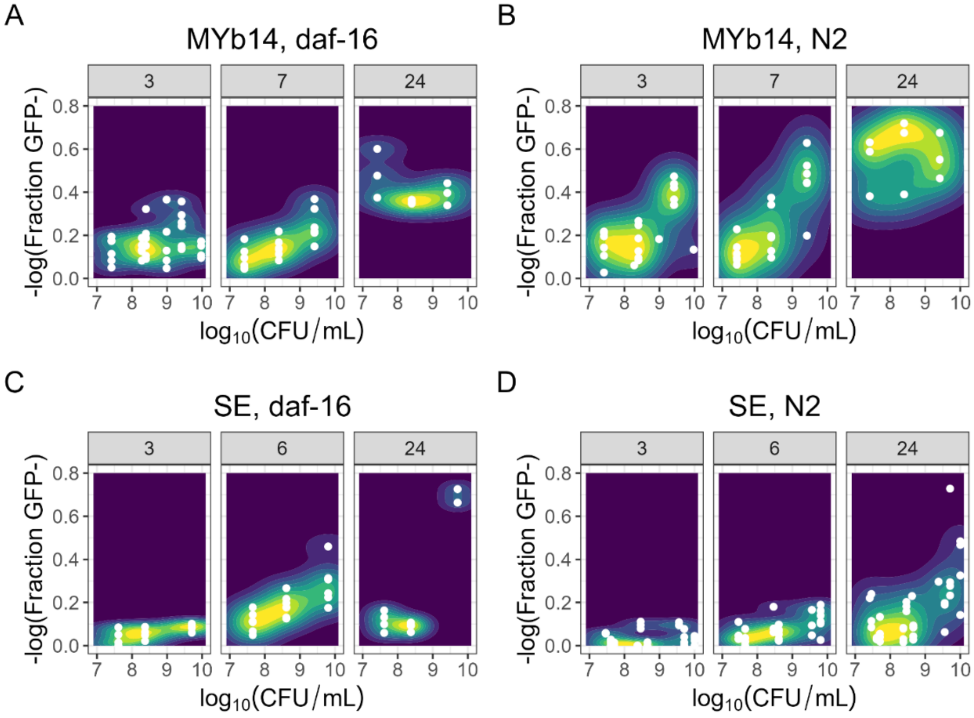


Figure S5. Kernel-smoothed distributions of susceptibility vs dose, with data sets split up by duration of exposure (time in hours, top bars). For MYb14-GFP, exposure durations of 2.5 and 4.5 hours were indistinguishable and were combined into a “3 hour” data set. For *S. enterica*, exposures of 6-7 hours were likewise combined into a “6 hour” data set. Color indicates relative density of the data; data points are shown in white.

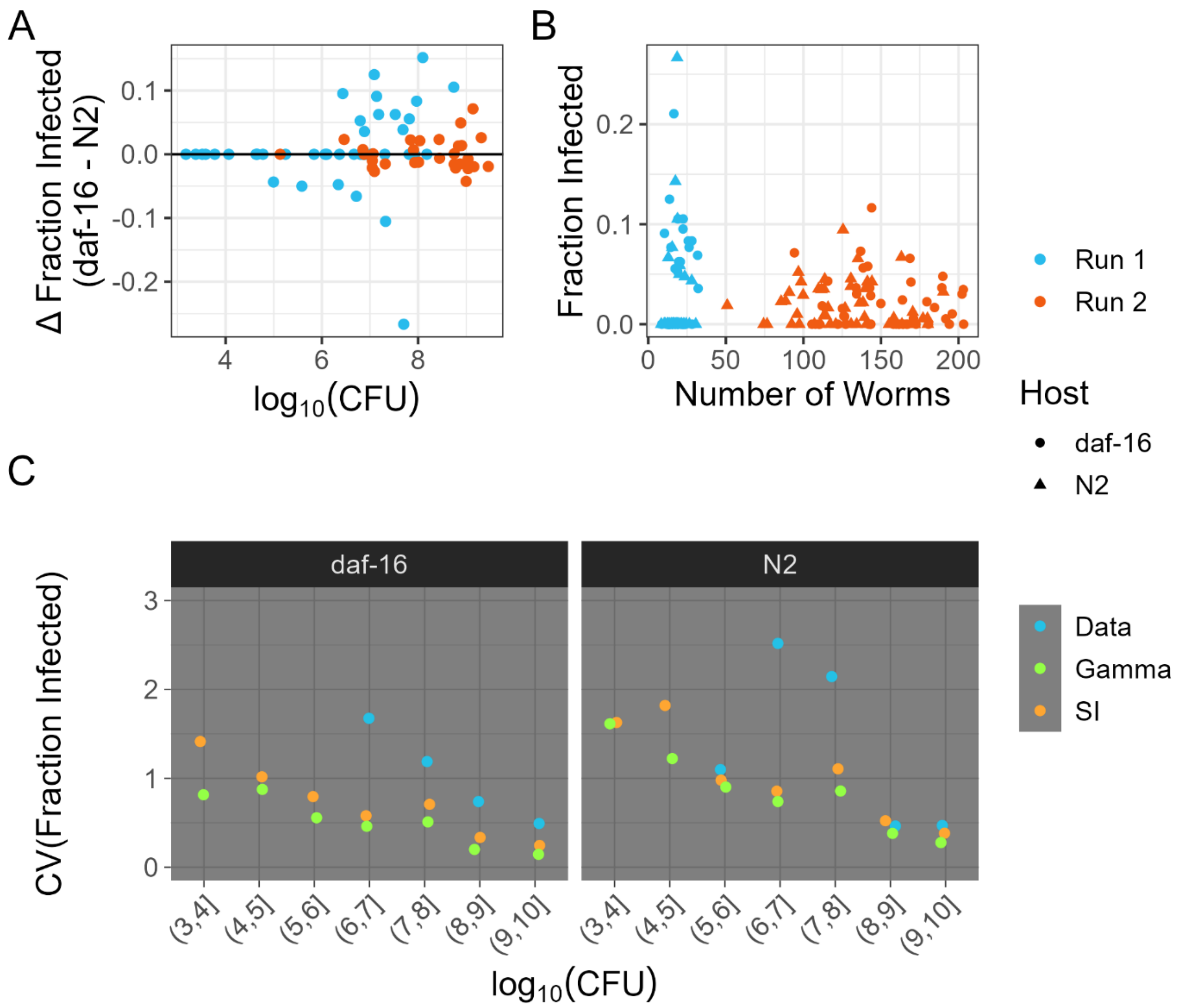


Figure S6. Trends in secondary infections. Original data are in **Figures 3+4**. (A) Difference in fraction infected in populations of N2 vs *daf-16* worms exposed to the same transmission event. Each point represents one pair of wells which received the same index case supernatant. Black solid line represents the median of the data. (B) Fraction of infected worms vs. total size of the susceptible population. (C) Coefficient of variation in fraction infected individuals across real (data) or simulated (SI/Gamma) susceptible populations, binned by bacterial density used for exposure. Colors indicate the susceptibility model used in simulations; experimental results (blue) are shown for comparison. (objects pSeSuscept_DiffInfect_bylogCFU, pSeSecondary_fracInfect_nworms, pSeSecondary_InfectVariation_summary)

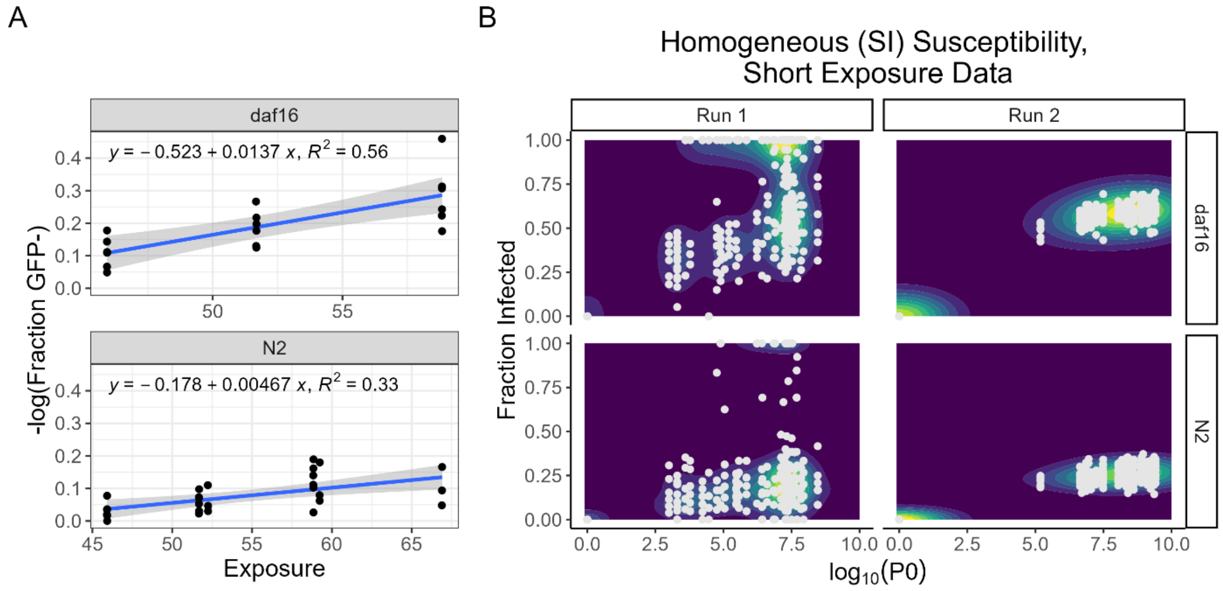


Figure S7. Predictions of the homogeneous model when fit to data for short exposure times. (A) Linear fits to 6-7 hour susceptibility data (subset of data from **Figure 3**). Grey bars are 95% CI. Note that these fits are against total exposure (log(CFU)*time) to allow use of the resulting parameters in time-dependent Gillespie simulations. (B) Simulation results using the models in (A). Simulated data points are shown in light grey. Contour map colors indicate density of simulated results (low=dark blue, high=yellow). (pSuscept_7h_LinearFits, psusceptOUT_SI_7h_contour)

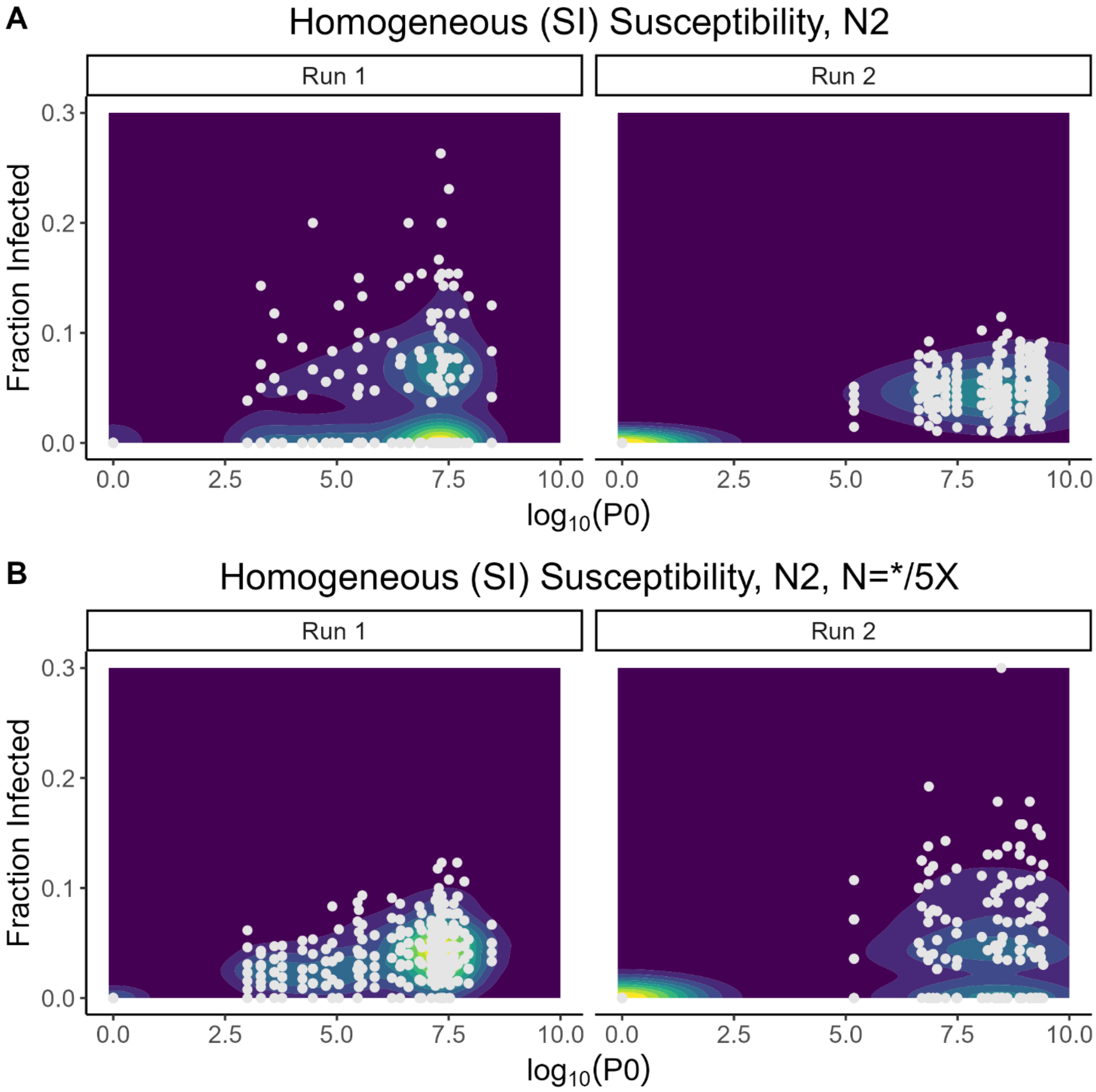


Figure S8 (psusceptOUT_SI_global_N2_POPSIZE_contour). Changing the number of worms per susceptible population alters predicted variation across simulated outbreaks. The original simulations (A) used the counts of worms per well taken from the data (**Figure 3**). Simulations with altered population sizes (B) were initiated with (run 1) 5X and (run 2) 0.2X susceptible population sizes in each well. Bacterial densities per well, and all other parameters, are the same in both sets of simulations. Ten simulation-based predictions of secondary infections were made for each combination of bacterial density and susceptible population.
